# Remodeling of ER-exit sites initiates a membrane supply pathway for autophagosome biogenesis

**DOI:** 10.1101/168518

**Authors:** Liang Ge, Min Zhang, Samuel J Kenny, Dawei Liu, Miharu Maeda, Kota Saito, Anandita Mathur, Ke Xu, Randy Schekman

## Abstract

Autophagosomes are double-membrane vesicles generated during autophagy. Biogenesis of the autophagosome requires membrane acquisition from intracellular compartments, the mechanisms of which are unclear. We previously found that a relocation of COPII machinery to the ER-Golgi intermediate compartment (ERGIC) generates ERGIC-derived COPII vesicles which serve as a membrane precursor for the lipidation of LC3, a key membrane component of the autophagosome. Here we employed super-resolution microscopy to show that starvation induces the enlargement of ER-exit sites (ERES) positive for the COPII activator, SEC12, and the remodeled ERES patches along the ERGIC. A SEC12 binding protein, CTAGE5, is required for the enlargement of ERES, SEC12 relocation to the ERGIC, and modulates autophagosome biogenesis. Moreover, FIP200, a subunit of the ULK protein kinase complex, facilitates the starvation-induced enlargement of ERES independent of the other subunits of this complex and associates via its C-terminal domain with SEC12. Our data indicate a pathway wherein FIP200 and CTAGE5 facilitate starvation-induced remodeling of the ERES, a prerequisite for the production of COPII vesicles budded from the ERGIC that contribute to autophagosome formation.

## Introduction

Autophagy is a fundamental catabolic process for bulk turnover of cytoplasmic components which plays essential roles in a broad range of physiological processes and pathological conditions, such as aging, cancer, neurodegeneration and metabolic syndromes [1,2]. Autophagy is usually activated by stress, such as starvation, during which multiple cascades of signaling pathways converge on a group of autophagic regulators, called autophagy-related gene (ATG) proteins [3,4]. These ATG proteins act hierarchically on the preautophagosomal structure/phagophore assembly site (PAS) located at a special region of the endoplasmic reticulum to assemble membranes from multiple locations into a cup-shaped membrane precursor called the phagophore. The phagophore engulfs a portion of the cytoplasm and seals at its extremity to form a double-membrane vesicle, termed the autophagosome. The autophagosome fuses with the endolysosome and becomes the autolysosome where the enclosed cytoplasmic contents are degraded by the hydrolases in the lysosome [5-9]. After degradation, the lysosomes are regenerated by pinching membranes off the autolysosome [10-13].

Autophagy requires multiple membrane remodeling events and protein catalysts whose activities are induced by starvation. One key event happens on the PAS where autophagosomal membrane precursors fuse, elongate, and close to form the autophagosome. Several autophagic protein machineries coordinate the formation of the PAS, including: the ULK protein kinase complex (FIP200/ULK1/ATG13/ATG101 complex), the phosphatidylinositol 3-kinase (PI3K) complex (ATG14/Beclin-1/P150/VPS34 complex), the ubiquitin-like phosphatidylethanolamine conjugation machinery (ATG7, ATG3 and ATG12-ATG5 conjugate in complex with ATG16), and the transmembrane protein ATG9 and downstream effectors, such as WIPIs and ATG2 [6,9]. Subsequently, the mature autophagosome fuses with the endolysosome followed by membrane tubulation and fission to regenerate new lysosomes [10]. The SNARE proteins Syntanxin-17, SNAP29 and VAMP8 as well as the HOPS complex participate in autophagosome-lysosome fusion [14-16], and autolysosome fission is mediated by clathrin and kinesin [12,17].

Molecular aspects of the initiation of phagophore assembly and the processes induced by starvation remain unclear. The endoplasmic reticulum (ER) is a major component of the endomembrane system. Emerging evidence indicates the involvement of ER and the associated compartments in autophagosome biogenesis. A subdomain of ER, possibly adjacent to the mitochondria, has been shown to act as a cradle for autophagosome biogenesis by forming a phosphatidylinositol 3-phosphate-enriched omegasome [18-21]. Multiple lines of investigation have suggested that the ER-Golgi intermediate compartment (ERGIC), a recycling station located between the ER and Golgi, is a membrane source for the autophagosome [22-27]. The ER-exit site (ERES), where COPII vesicle formation initiates the traffic of secretory cargo from the ER is associated with the PAS in yeast [28-30]. In addition, the COPII components as well as TRAPPIII, a proposed molecular machinery for tethering COPII vesicles and a secretory pathway regulator, are required for autophagy in both yeast and mammalian cells [31-34]. The molecular links between these steps remain to be elucidated.

In previous work we reported a mechanism of membrane remodeling of the ERGIC wherein a starvation-induced relocation of COPII machinery from the ERES to the ERGIC generates ERGIC-derived COPII vesicles to serve as membrane templates for LC3 lipidation, a key step of autophagosome biogenesis [25]. The assembly of COPII vesicles on the ERGIC is achieved by a relocation of SEC12, the activator of COPII assembly, from the ERES to the ERGIC [25]. We sought to understand how SEC12 is mobilized from the ER during starvation. Here we show that a starvation-induced remodeling of the ERES positive for SEC12 (SEC12-ERES) facilitates the relocation of SEC12 to the ERGIC. Three-dimensional (3D) super-resolution microscopy indicates that the SEC12-ERES structure is enlarged and surrounds the ERGIC, leading to the relocation of SEC12 to the ERGIC, possibly to trigger ERGIC-COPII vesicle assembly. Depletion of CTAGE5 and FIP200, two proteins associated with SEC12 which are required for the concentration of SEC12 on the ERES, abolishes the starvation-induced enlargement of SEC12-ERES, the relocation of SEC12 to the ERGIC, and LC3 lipidation. Thus we suggest that a SEC12 protein complex is the target of control that mobilizes SEC12 from the ERES to the ERGIC as an early event in starvation-induced autophagy.

## Results

### Starvation-induced remodeling of SEC12-ERES facilitates SEC12 relocation to the ERGIC

SEC12 is a type II transmembrane protein that normally localizes to the ERES where it serves to activate nucleotide exchange on SAR1 to initiate COPII assembly [28]. To investigate how starvation leads to the relocation of COPII from the ERES to the ERGIC, we analyzed endogenous SEC12 and ERGIC53 (a marker of the ERGIC) by confocal microscopy (Fig 1A). Under nutrient-rich conditions, SEC12 was localized in punctate compartments (ERES), some of which were associated with the ERGIC (Fig EV1A, Ctr). Upon starvation, a fraction of the SEC12 compartments increased in size and colocalized with the ERGIC (Fig EV1A, ST). We counted the fraction of SEC12 puncta above 0.1 μm^2^ over the total and found a ^~^1.5 fold increase after starvation (Fig EV1B). (see Methods in detail).

**Figure 1.**
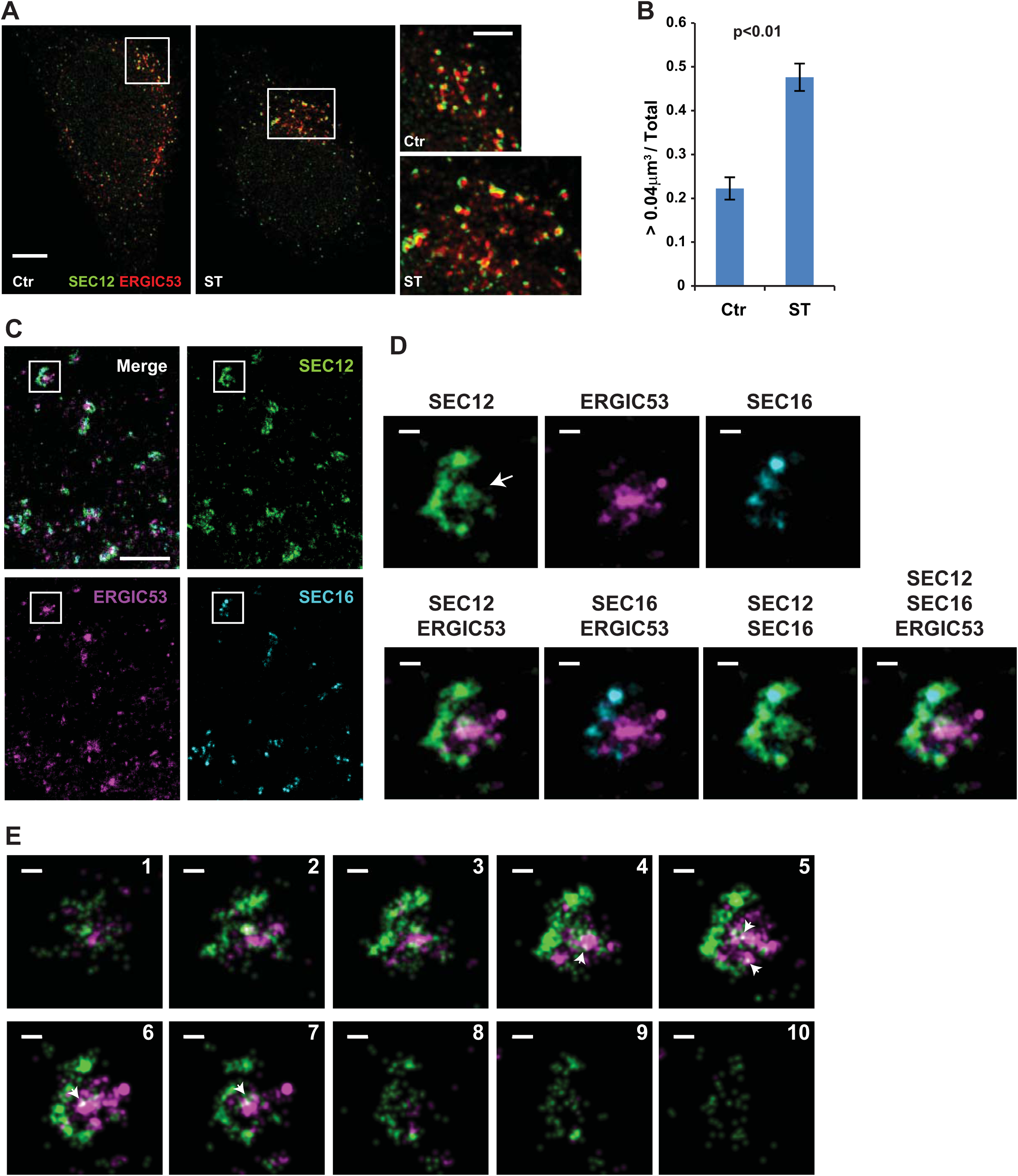
Remodeling of SEC12-positive compartment upon starvation. (A) Hela cells were incubated in nutrient-rich medium (Ctr) or starved in EBSS (ST) for 1 h. Structured illumination microscopy (SIM) was performed to analyze the structure and relation of SEC12 and ERGIC compartments. Bars are 5 μm (original image) and 2 μm (zoomed in image) respectively. (B) Quantification of the fraction of SEC12 compartments larger than 0.04 μm^3^ in volume analyzed in (A). Error bars represent standard deviations. P value was obtained from Two-tailed T test. Five experiments were performed for the statistics. (C) Hela cells were starved in EBSS for 1 h. Immunofluorescence and 3D-STORM were performed to resolve the structure of SEC12 and ERGIC compartments. Bar: 1 μm (D) Zoomed in images of the boxed areas in (D). Arrows indicate the SEC12 localized on the ERGIC area. Bar: 100 nm (E) Virtual Z-sections (50 nm thickness and 40 nm per step) of (D) with SEC12 and ERGIC. Of the total, 10 of 13 sections are shown. Arrows indicate the ERGIC-localized SEC12 (D). Bar: 100 nm

**Expanded View Figure 1.**
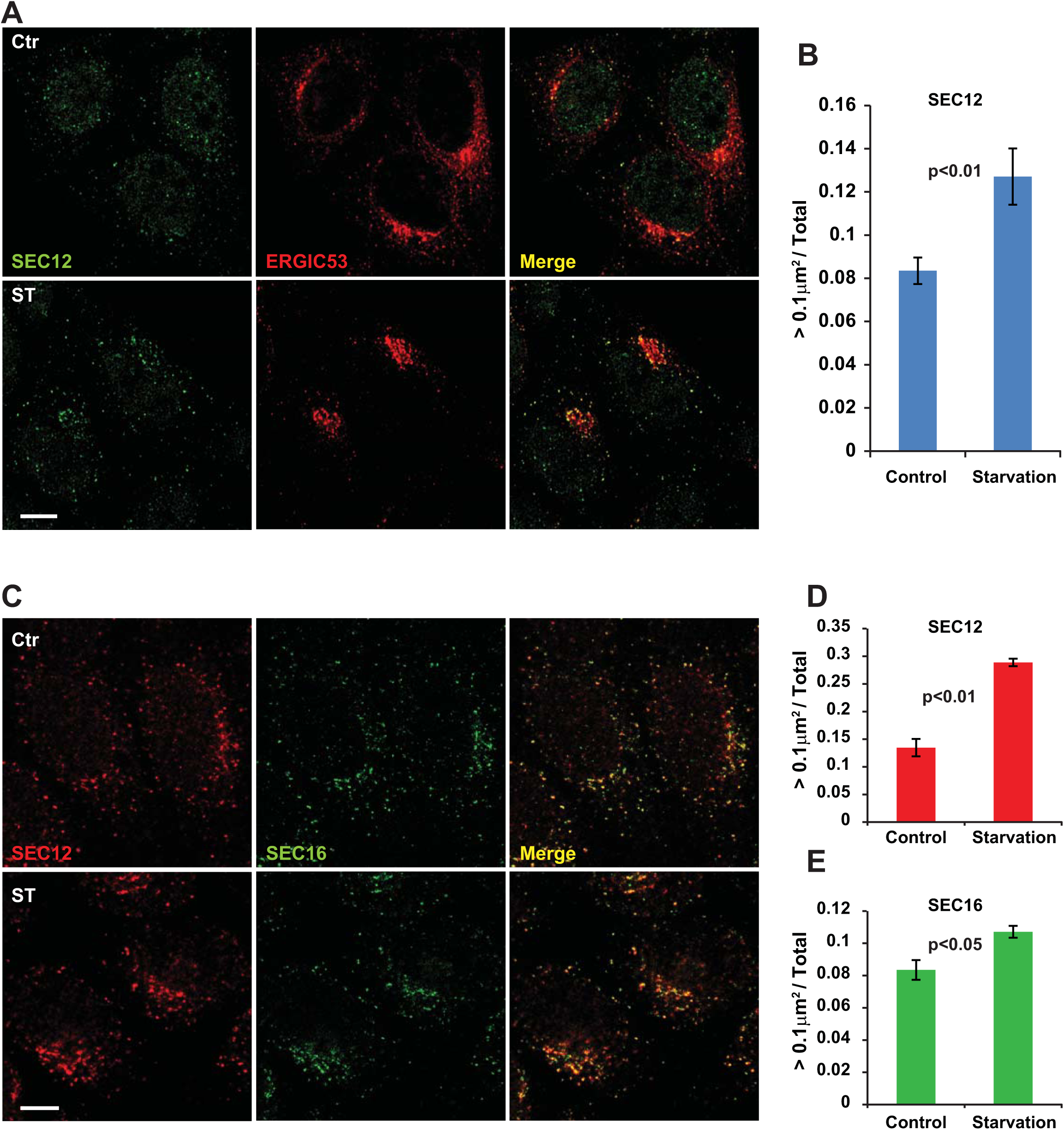
Enlargement of SEC12-positive compartment upon starvation. (A) Hela cells were incubated in nutrient-rich medium (Ctr) or starved in EBSS (ST) for 1 h. Immunofluorescence and confocal microscopy were performed to visualize SEC12 and ERGIC53. Bar: 10 μm (B) Quantification of the fraction of SEC12 compartments larger than 0.1 μm^2^ in area analyzed in (A). Error bars represent standard deviations. P value was obtained from Two-tailed T test. Five experiments (50-100 cell/experiment) were performed for the statistics. (C) Hela cells were incubated in nutrient-rich medium (Ctr) or starved in EBSS (ST) for 1 h. Immunofluorescence and confocal microscopy were performed to visualize SEC12 and SEC16 compartments. Bar: 10 μm (D) Quantification of the percentage of SEC12 compartments larger than 0.1 μm^2^ in area analyzed in (C). Error bars represent standard deviations. P value was obtained from Two-tailed T test. Five experiments (50-100 cell/experiment) were performed for the statistics. (E) Quantification of the percentage of SEC16 compartments larger than 0.1 μm^2^ in area analyzed in (C). Error bars represent standard deviations. P value was obtained from Two-tailed T test. Five experiments (50-100 cell/experiment) were performed for the statistics.

To determine if the starvation-induced effect on the ERES was SEC12 specific, we analyzed SEC16, a scaffold protein required for the proper organization of the ERES [35,36]. The fraction of SEC16 puncta bigger than 0.1 μm^2^ also increased upon starvation (^~^1.2 fold increase) albeit less dramatically than that of SEC12 puncta (^~^2 fold increase, Fig EV1C-E), suggesting that the overall ERES is altered by starvation with a greater effect on the SEC12-positive domains. For simplicity, we refer to the SEC12-positive domain of the ERES as SEC12-ERES.

We previously found that the autophagic PI3K is required for the relocation of COPII machinery to the ERGIC and for the generation of ERGIC-COPII vesicles. We reexamined the SEC12-ERES in cells starved in the presence of wortmannin or 3-methyladenine, two inhibitors of PI3K [37] (Fig EV2A, B). We also employed three VPS34-specific inhibitors PIK-III [38]: SAR405[39], and VPS34-IN1[40] (Fig EV2C, D). The enlargement of SEC12-ERES was not inhibited but instead slightly enhanced by several of the PI3K inhibitors (Fig EV2), suggesting that PI3K may act in a step downstream of the remodeling of SEC12-ERES.

**Expanded View Figure 2.**
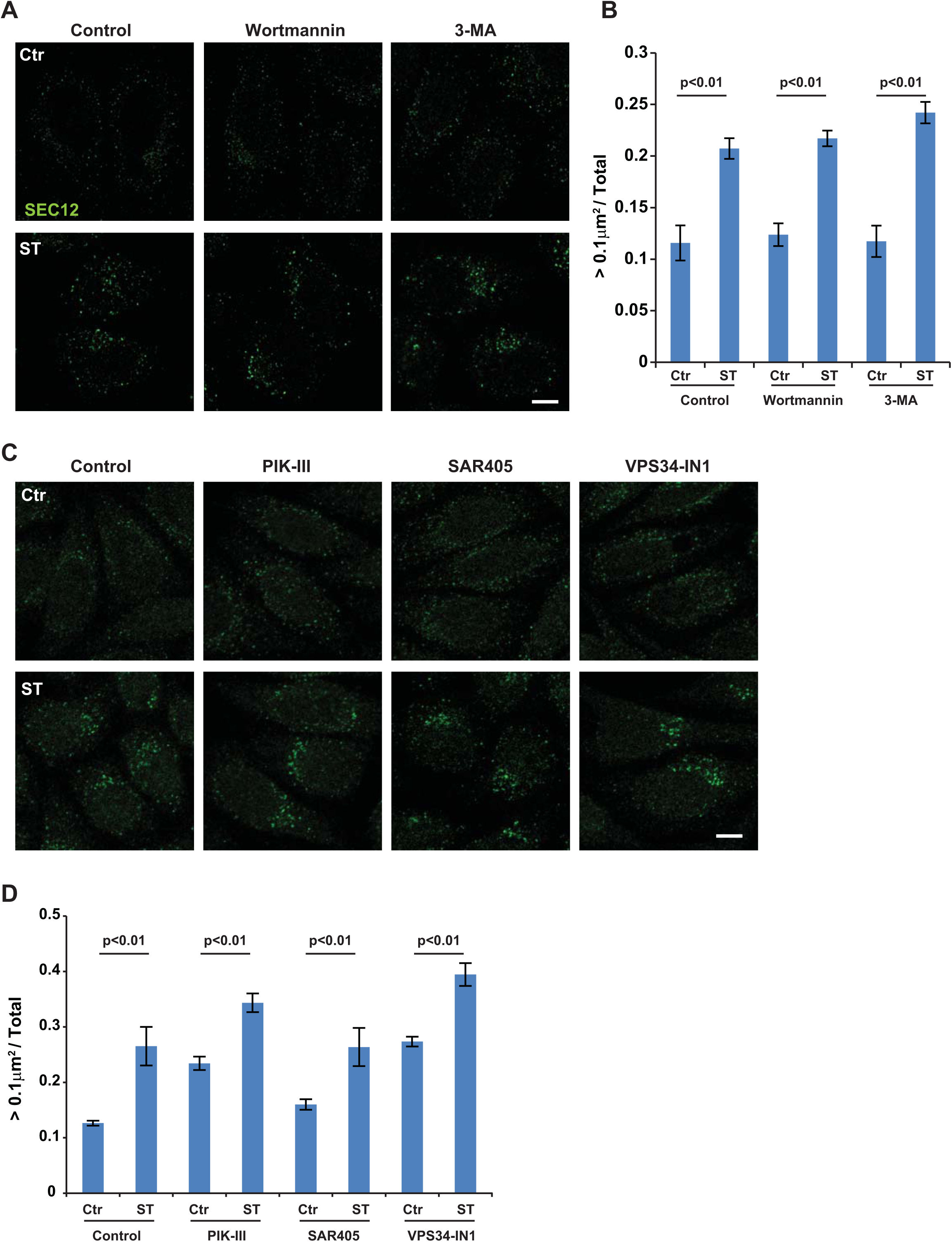
PI3K is not required for the starvation-induced enlargement of SEC12 compartment. (A) Hela cells were incubated in nutrient-rich medium (Ctr) or starved in EBSS (ST) with control, wortmannin (10nM) or 3-methyladenine (3-MA, 10mM) for 1 h. Immunofluorescence and confocal microscopy were performed to visualize SEC12 compartments. Bar: 10 μm (B) Quantification of the fraction of SEC12 compartments larger than 0.1 μm^2^ in area analyzed in (A). Error bars represent standard deviations. P value was obtained from Two-tailed T test. Five experiments (50-100 cell/experiment) were performed for the statistics. (C) Hela cells were incubated in nutrient-rich medium (Ctr) or starved in EBSS (ST) with control, PIK-III, SAR405 or VPS34-IN1 (20 μM each) for 1 h. Immunofluorescence and confocal microscopy were performed to visualize SEC12 compartments. Bar: 10 μm (D) Quantification of the fraction of SEC12 compartments larger than 0.1 μm^2^ in area analyzed in (C). Error bars represent standard deviations. P value was obtained from Two-tailed T test. Five experiments (50-100 cell/experiment) were performed for the statistics.

To further define the structure of SEC12-ERES, we performed structured illumination microscopy (Fig 1A). We observed punctate SEC12-ERES structures close to the ERGIC under nutrient-rich conditions that elongated and some formed cup-shaped compartments surrounding the ERGIC upon starvation (Fig 1A). On average, around ten elongated SEC12-ERES compartments were observed in each cell after starvation versus fewer than two before starvation (data not shown). The increased volume of the SEC12-ERES was quantified from Z-section stacks. As shown in Fig 1B, the fraction of SEC12-ERES larger than 0.04 μm^3^ (corresponding to the area of ^~^0.1 μm^2^) increased about twofold after starvation, confirming the enlargement of SEC12-ERES in 3D space.

To further resolve the structural details of the SEC12-ERES with the surrounded ERGIC upon starvation, we employed 3D stochastic optical reconstruction microscopy (3D-STORM) to achieve ^~^20 nm resolution (Fig 1C-E and Appendix Fig S1 & S2) [41,42]. Visual Z-stack images indicated that elongated SEC12-ERES (green) patched along the ERGIC (magenta), which documented a close association between the two compartments (Fig 1C, D). SEC16 (cyan) was condensed in distinct regions on the SEC12-ERES, consistent with a role as a scaffold for the ERES (Fig 1D). A diffusely stained area of SEC12, possibly overlapped with the ERGIC, also appeared close to some of the SEC12-ERES (Fig 1D, arrow pointed). This is reminiscent of the SEC12 relocation to the ERGIC reported in our previous work [25]. To confirm the localization of SEC12 on the ERGIC upon starvation, we examined Z-sections of the structure shown in Fig 1E and Appendix Fig S1 & S2. Overlap between SEC12 and the ERGIC was observed in some of the sections (Fig 1E, sections 4-7, arrow pointed, Appendix Fig S1A and Appendix Fig S2C, D, arrow pointed), consistent with the proposed localization of SEC12 on the ERGIC. The data were consistent with our previous findings using biochemical fractionation in which SEC12 was shown to localize on the ERGIC isolated from starved cells. Notably, SEC16 overlapped with part of the SEC12-ERES but not the ERGIC (Fig 1 and Appendix Fig S1 & S2), indicating it was not relocated to the ERGIC upon starvation.

These data indicated that starvation induced the remodeling of SEC12-ERES by increasing the compartment size and its association with the ERGIC. This association may permit SEC12 to be relocated from the ERES to the ERGIC, possibly as a step to trigger COPII assembly in order to generate LC3 lipidation-active precursor vesicles.

### CTAGE5 is required for the remodeling of SEC12-ERES

To examine the molecular requirements involved in the remodeling of SEC12-ERES, we performed co-immunoprecipitation coupled with mass spectrometry to survey SEC12-interacting proteins. CTAGE5 was identified as a protein that strongly associated with SEC12 (data not shown). Because CTAGE5 has been shown to be required for the concentration of SEC12 on the ERES [43], we hypothesized that CTAGE5 was involved in the remodeling of SEC12-ERES. We confirmed the association between CTAGE5 and SEC12 in a co-immunoprecipitation (co-IP) experiment (Fig EV3A). In addition, we found that the protein complex of CTAGE5 and SEC12 did not substantially change in nutrient rich and starvation conditions (Fig EV3A). Immunofluorescence analysis of endogenous CTAGE5 and SEC12 showed that they are colocalized in nutrient-rich conditions (Fig 2A and Fig EV3B, C). Starvation increased the size of CTAGE5 puncta which remained overlapped with SEC12 (Fig 2A-C and Fig EV3B, C). This coincidence suggested a role for CTAGE5 in the starvation-dependent remodeling of the SEC12-ERES.

**Expanded View Figure 3.**
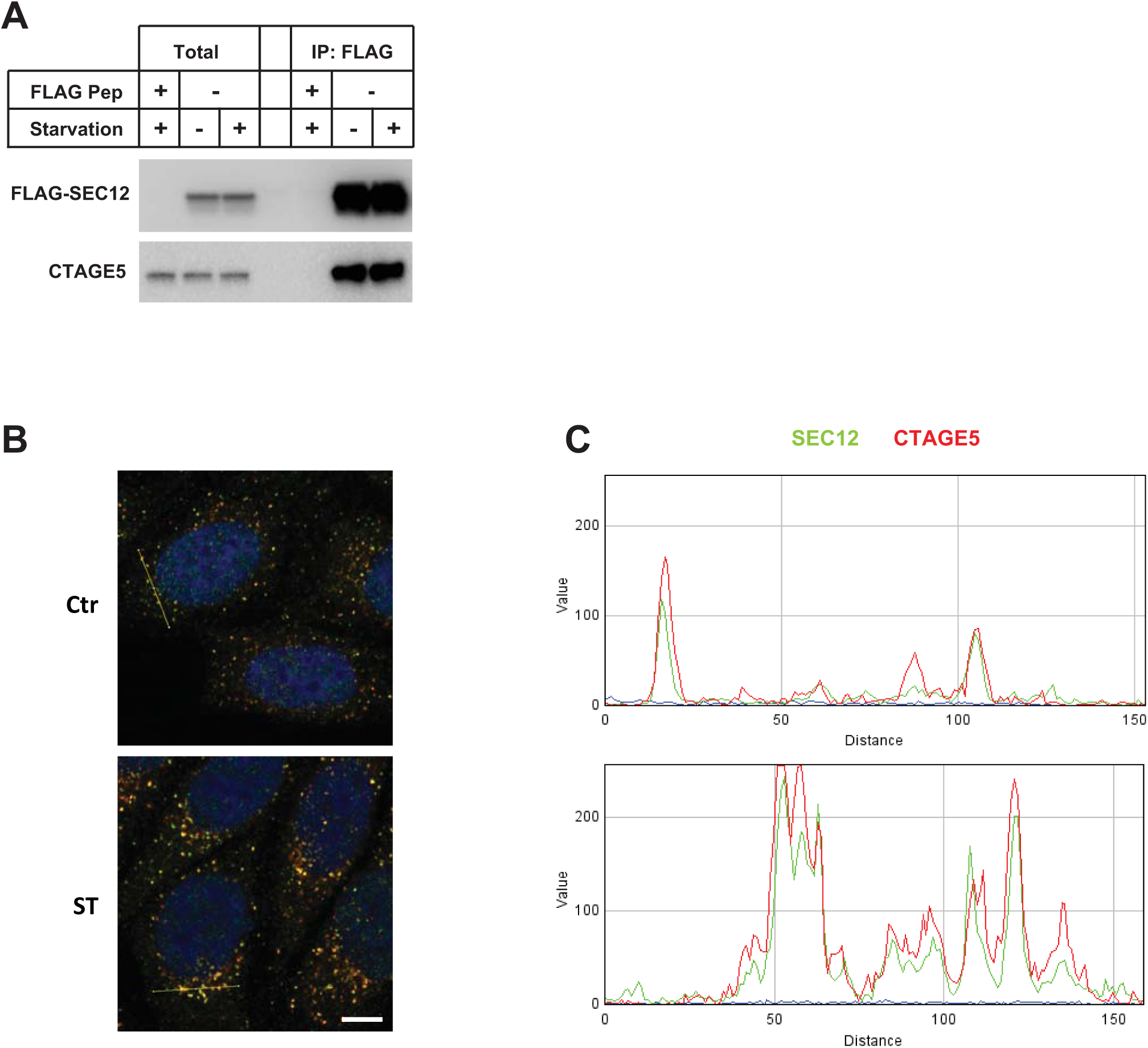
Analysis of SEC12 and CTAGE5 colocalization and requirement of CTAGE5 for LC3 lipidation. (A) HEK293T cells were transfected with control plasmids or plasmids encoding FLAG-SEC12. After 24 h, the cells were incubated in nutrient-rich medium or starved in EBSS for 1 h. Immunoprecipitation of FLAG-SEC12 was performed and the levels of indicated proteins from indicated fractions were determined by immunoblot. (B, C) RGB profile plots showing the colocalization of SEC12 (green) and CTAGE5 (red) through the lines drawn on the merged images shown in Figure 2A.

**Figure 2.**
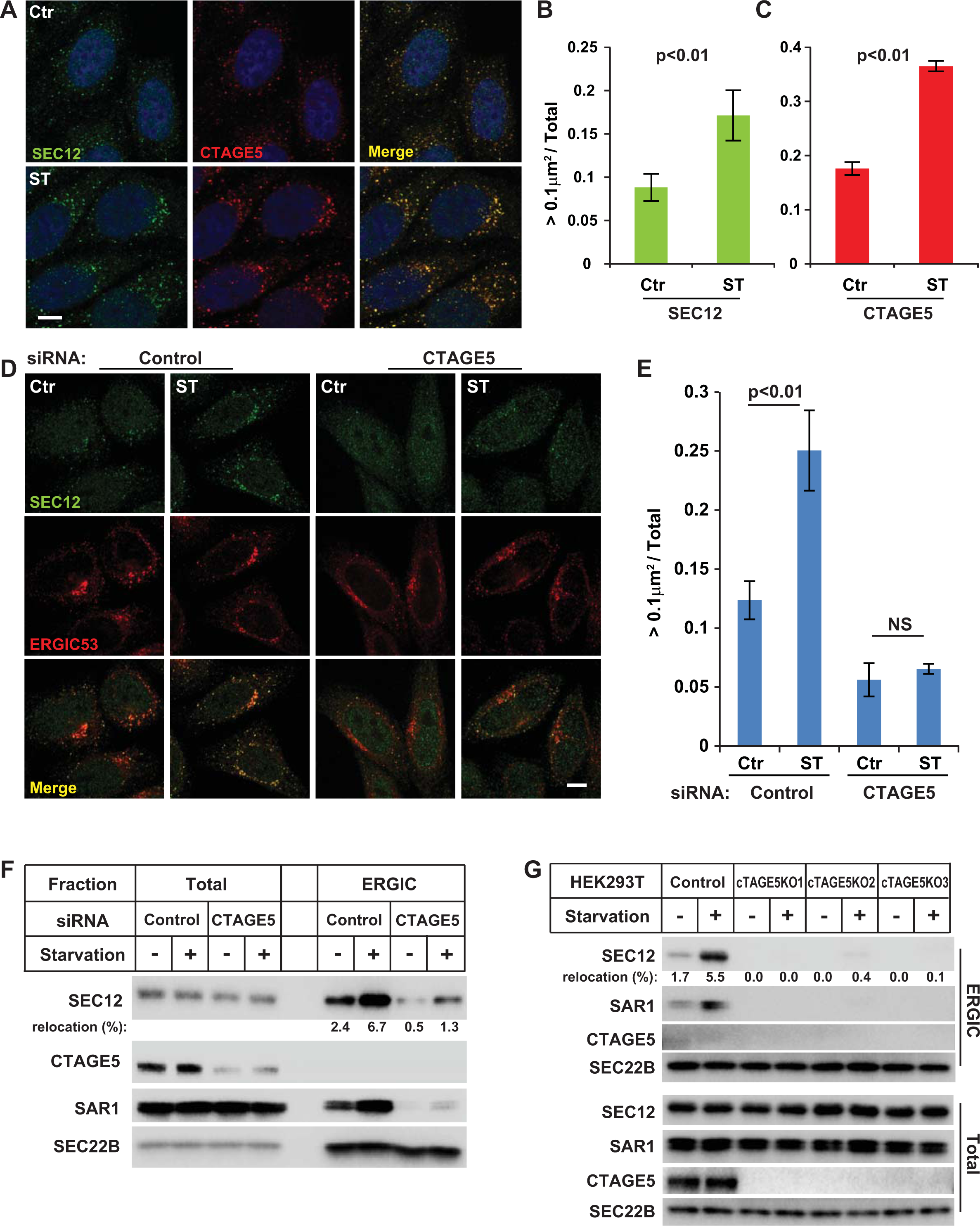
Involvement of CTAGE5 in the enlargement of SEC12 compartment and relocation of SEC12 to the ERGIC upon starvation. (A) Hela cells were incubated in nutrient-rich medium (Ctr) or starved in EBSS (ST) for 1 h. Immunofluorescence and confocal microscopy were performed to visualize SEC12 and CTAGE5. Bar: 10 μm (B, C) Quantification of the fraction of SEC12 or CTAGE5 compartments larger than 0.1 μm^2^ in area analyzed in (A). Error bars represent standard deviations. P value was obtained from Two-tailed T test. Five experiments (50-100 cell/experiment) were performed for the statistics. (D) Hela cells were transfected with control and siRNAs against cTAGE5. After 72 h, the cells were incubated in nutrient-rich medium (Ctr) or starved in EBSS (ST) for 1 h Immunofluorescence and confocal microscopy were performed to visualize SEC12 and ERGIC53. Bar: 10 μm (E) Quantification of the fraction of SEC12 or CTAGE5 compartments larger than 0.1 μm^2^ in area analyzed in (D). Error bars represent standard deviations. P value was obtained from Two-tailed T test. Five experiments (50-100 cell/experiment) were performed for the statistics. (F) HEK293T cells were transfected with control and siRNAs against cTAGE5. After 72 h, the cells were incubated in nutrient-rich medium or starved in EBSS for 1 h. The cell lysates and the ERGIC membrane fractions were analyzed by immunoblot to determine the levels of indicated proteins. Quantification shows the percentage of SEC12 relocation to the ERGIC under the indicated conditions. (G) cTAGE5 knockout (KO) cell lines were established using CRISPR/Cas9 in HEK293T cells. Control cells as well as three KO cell lines were incubated in nutrient-rich medium or starved in EBSS for 1 h. The cell lysates and the ERGIC membrane fractions were analyzed by immunoblot to determine the levels of indicated proteins. Quantification shows the percentage of SEC12 relocation to the ERGIC under the indicated conditions.

To test the role of CTAGE5 in the maintenance of SEC12-ERES, we knocked down cTAGE5 expression by siRNA-mediated gene silencing. As shown in Fig 2D, the size of SEC12-ERES in control cells was increased upon starvation, whereas SEC12 was dispersed under both nutrient-rich and starved conditions in cTAGE5 knockdown (KD) cells (Fig 2D). Quantification showed that the fraction of SEC12 puncta (>0.1μm^2^) in cTAGE5 KD cells was reduced (^~^50% decrease versus that of control cells under nutrient-rich condition) and not increased by starvation (Fig 2E). The dispersed localization of SEC12 caused by cTAGE5 KD is consistent with the recent study showing the requirement of CTAGE5 for the enrichment of SEC12 on the ERES [43]. In addition, the effect of CTAGE5 was specific for positioning SEC12 on the ERES because it was reported that cTAGE5 KD does not affect the localization of the ERES scaffold protein SEC16 or general ER-Golgi trafficking [43]. Therefore, CTAGE5 is required for the concentrated localization of SEC12 on the ERES, as well as the starvation-induced enlargement of SEC12-ERES.

To determine the role of CTAGE5 in the relocation of SEC12 to the ERGIC upon starvation, we performed cTAGE5 KD and isolated the ERGIC fraction (Fig 2F). In control cells, the localization of SEC12 on the ERGIC was increased upon starvation (^~^3 fold increase) which was consistent with our previous finding (Fig 2F) [25]. Although CTAGE5 associated with SEC12 in unfractionated cell lysates (Fig EV3A), CTAGE5 was not localized on the ERGIC in starved cells (Fig 2F). KD of cTAGE5 inhibited the localization of SEC12 on the ERGIC (Fig 2F). A small fraction (^~^20% of the wt control) of SEC12 relocated to the ERGIC upon starvation in cTAGE5 KD cells, possibly because of residual CTAGE5 in KD cells (Fig 2F). Complete depletion of CTAGE5 by CRISPR/CAS9 genomic editing abolished the localization of SEC12 on the ERGIC in all 3 knockout (KO) cell lines (Fig 2G).

Therefore, the data demonstrated that CTAGE5 is associated with SEC12, possibly only on the ERES, and is required for the enrichment of SEC12 on the ERES as well as its relocation to the ERGIC upon starvation.

### CTAGE5 modulates LC3 lipidation and autophagosome biogenesis under starvation

We next sought to determine the role of CTAGE5 in autophagosome biogenesis. In control cells, we observed starvation-increased lipidation of LC3 (LC3-II), a marker of the autophagosome (Fig 3A) [44]. Addition of Bafilomycin A1 to block lysosomal acidification and protein degradation caused lipidated LC3 to accumulate to a higher extent in starved cells (Fig 3A). The level of LC3-II was decreased in cTAGE5 KD (^~^ 50% decrease) and was reduced even more in cTAGE5 KO cells (^~^ 75% decrease, Fig 3A, B).

**Figure 3.**
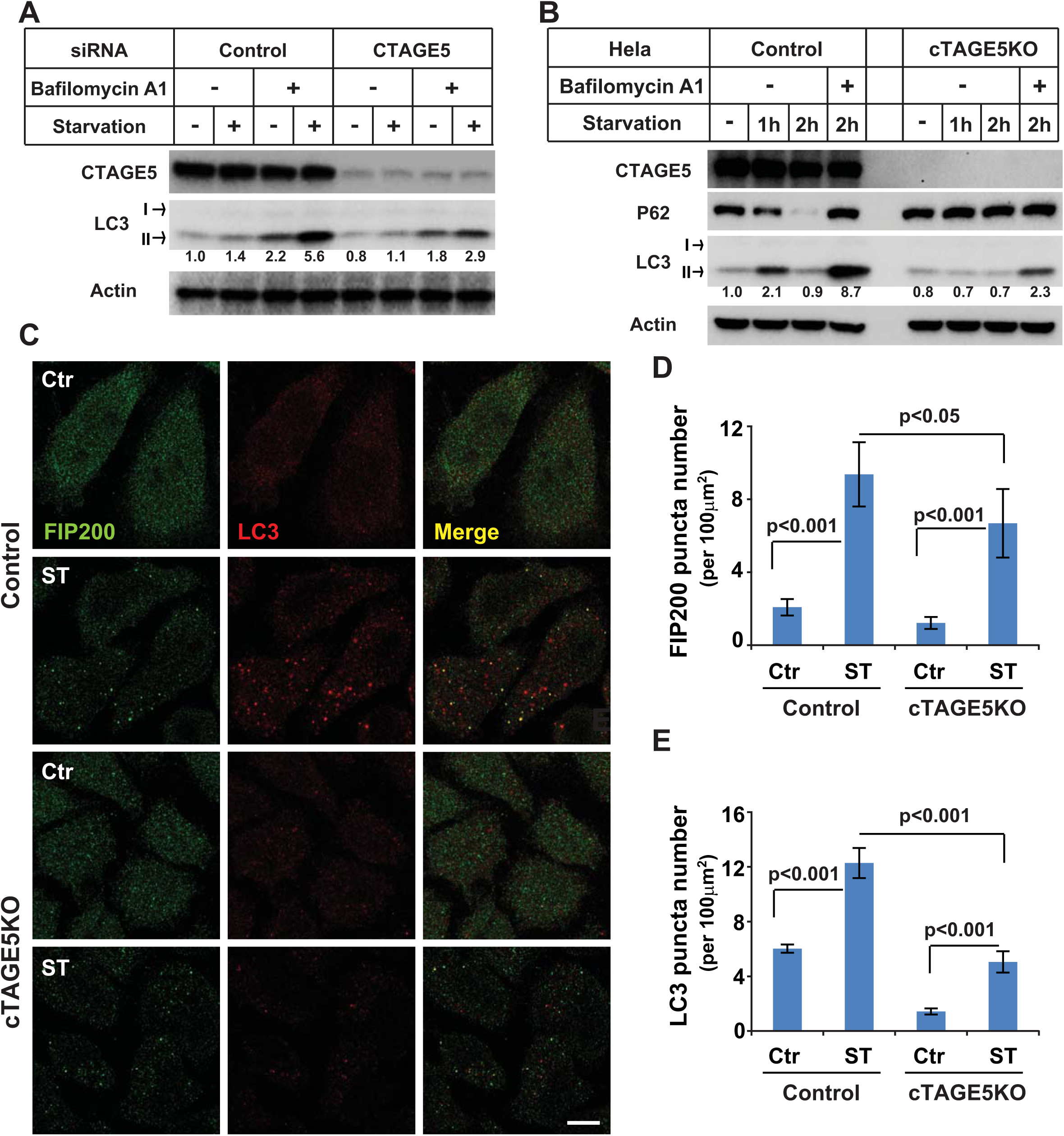
CTAGE5 modulates autophagosome biogenesis. (A) Hela cells were transfected with control and siRNAs against cTAGE5. After 72 h, the cells were incubated in nutrient-rich medium or starved in EBSS in the absence or presence of 500 nM Bafilomycin A1 for 1 h. Immunoblots was performed to determine the levels of indicated proteins. Quantification shows the relative level of LC3 lipidation to the control (control siRNA and before starvation in the absence of Bafilomycin A1). (B) Control and cTAGE5 knockout (KO) cells were incubated in nutrient-rich medium or starved in EBSS for 1 h and 2h in the absence or presence of Bafilomycin A1. Immunoblot was performed to determine the levels of indicated proteins. Quantification shows the relative level of LC3 lipidation to the control (control cell and before starvation). (C) Control cells and cTAGE5 knockout (KO) cells shown in (B) were incubated in nutrient-rich medium or starved in EBSS for 1 h. Immunofluorescence and confocal microscopy were performed to visualize FIP200 and LC3. Bar: 10 μm (D, E) Quantification of the FIP200 (D) and LC3 (E) puncta number analyzed in (C). Error bars represent standard deviations. P value was obtained from Two-tailed T test. Five experiments (50-100 cell/experiment) were performed for the statistics.

For autophagic activity, we determined the turnover of P62/SQSTM1, a cargo adaptor and substrate for autophagy (Fig 3B) [37]. In control cells, P62 was partially degraded by the lysosomal pathway after a 2 h-starvation Fig 4B). The degradation of P62 was compromised in starved cells treated with Bafilomycin A1 and in the cTAGE5 KO cell lines (Fig 3B). We also analyzed the fluctuation of LC3 lipidation as an indicator of autophagosome biogenesis and turnover (Fig 3B) [37]. In control cells, LC3 lipidation increased (1h) and then decreased (2h) during starvation (Fig 3B), consistent with autophagic progression. Bafilomycin A1 blocked LC3 turnover and led to its accumulation (Fig 3B). Again, the level of LC3 lipidation was decreased and the fluctuation was compromised in the cTAGE5 KO cell (Fig 3B).

**Figure 4.**
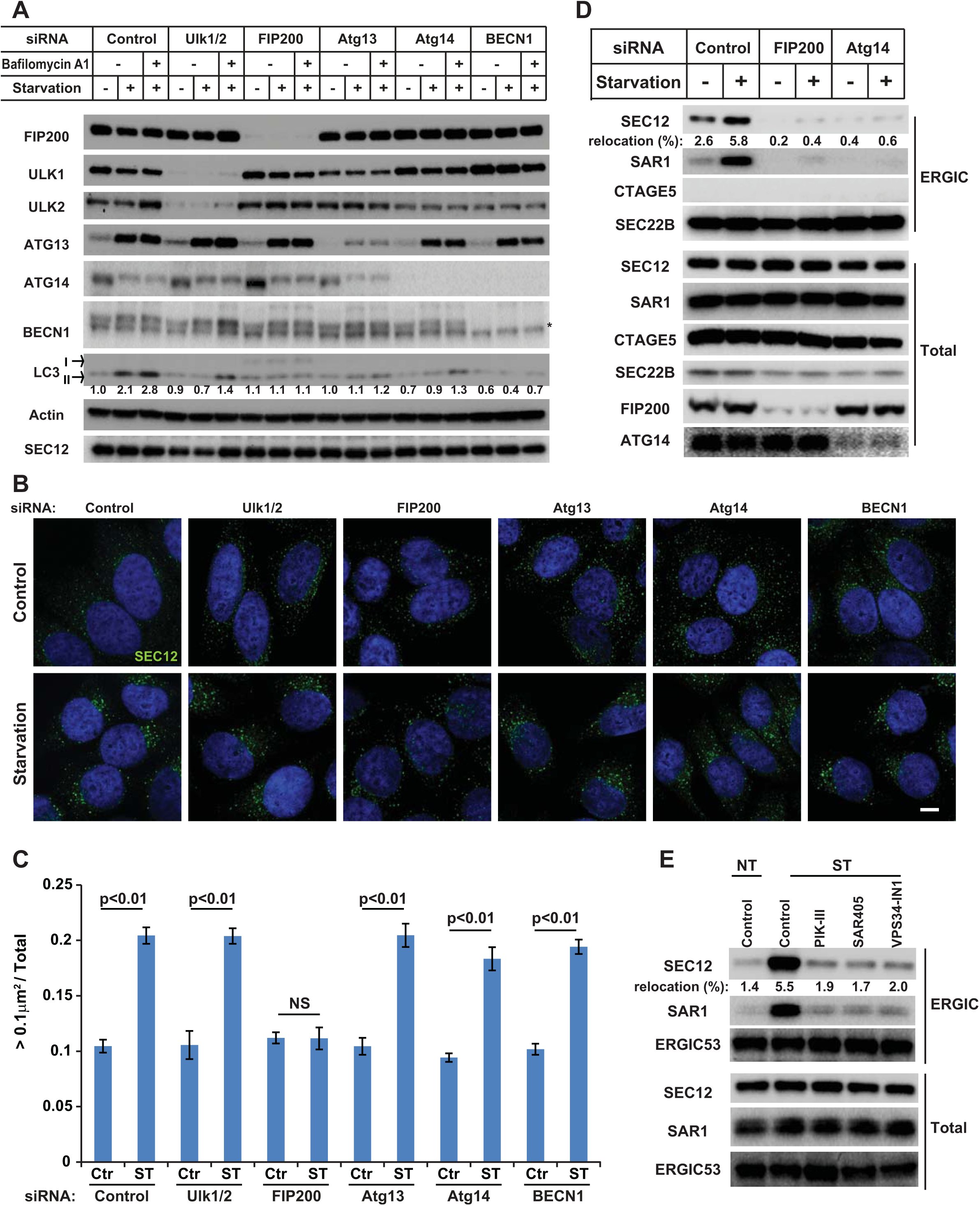
Requirement of FIP200 in the enlargement of SEC12-ERES compartment and the relocation of SEC12 to the ERGIC upon starvation. (A) Hela cells were transfected with control or siRNAs against the indicated autophagic factors. After 72 h, the cells were incubated in nutrient-rich medium or starved in EBSS for 1h in the absence or presence of Bafilomycin A1. Immunoblot was performed to determine the levels of indicated proteins. Quantification shows the relative level of LC3 lipidation to the control (control siRNA and before starvation in the absence of Bafilomycin A1). (B) Hela cells transfected with the indicated siRNAs were incubated in nutrient-rich medium (Control) or starved in EBSS (Starvation) for 1h. Immunofluorescence and confocal microscopy were performed to visualize SEC12. Bar: 10 μm (C) Quantification of the fraction of SEC12 compartments larger than 0.1 μm^2^ in area analyzed in (A). Error bars represent standard deviations. P value was obtained from Two-tailed T test. Five experiments (50-100 cell/experiment) were performed for the statistics. (D) HEK293T cells transfected with indicated siRNAs were incubated in nutrient-rich medium or starved in EBSS for 1h. Immunoblot was performed to determine the levels of indicated proteins from the total cell lysates and the ERGIC fractions. Quantification shows the percentage of SEC12 relocation to the ERGIC under the indicated conditions. (E) HEK293T cells were incubated in nutrient-rich medium or starved in EBSS without or with the indicated PI3K inhibitors for 1h.Immunoblot was performed to determine the levels of indicated proteins from the total cell lysates and the ERGIC fractions. Quantification shows the percentage of SEC12 relocation to the ERGIC under the indicated conditions.

We compared autophagosome biogenesis in control and cTAGE5 KO cells by fluorescence microscopy (Fig 3C). Membrane bound LC3-II served as an indicator of autophagosome biogenesis [44]. FIP200, an earlier marker of the autophagosome, was also used to mark the localization of the PAS [45]. Starvation-enhanced autophagosome biogenesis was inhibited in cTAGE5 KO cells (Fig 3C), whereas puncta marked by FIP200 were only moderately affected (Fig 3C). LC3 puncta decreased ^~^60% whereas FIP200 puncta decreased ^~^30% in starved cTAGE5 KO cells (Fig3D, E). Therefore, the data indicated that the CTAGE5-dependent remodeling of SEC12-ERES facilitated LC3 lipidation in multiple steps of autophagosome biogenesis with some selectivity for the LC3 lipidation step.

### FIP200 is required for the starvation-induced remodeling of SEC12-ERES

Recent studies showed that SEC12 forms a protein complex with CTAGE5 and TANGO1, which is essential for SEC12 concentration at the ERES and collagen VII secretion [43]. One possibility is that starvation may further enhance the association between SEC12 and CTAGE5 to form a bigger protein complex which leads to the enlargement of SEC12-ERES. We analyzed the size of the SEC12-CTAGE5-TANGO1 complex by Blue Native-PAGE as described before [46]. We found that the size of the protein complex is not altered by starvation or PI3K inhibition (Fig EV4), nor is the association between SEC12 and CTAGE5 in a co-IP experiment (Fig EV3A and Fig.5 described later). Therefore, mechanisms other than the SEC12-CTAGE5 association account for starvation-induced remodeling of SEC12-ERES.

**Expanded View Figure 4.**
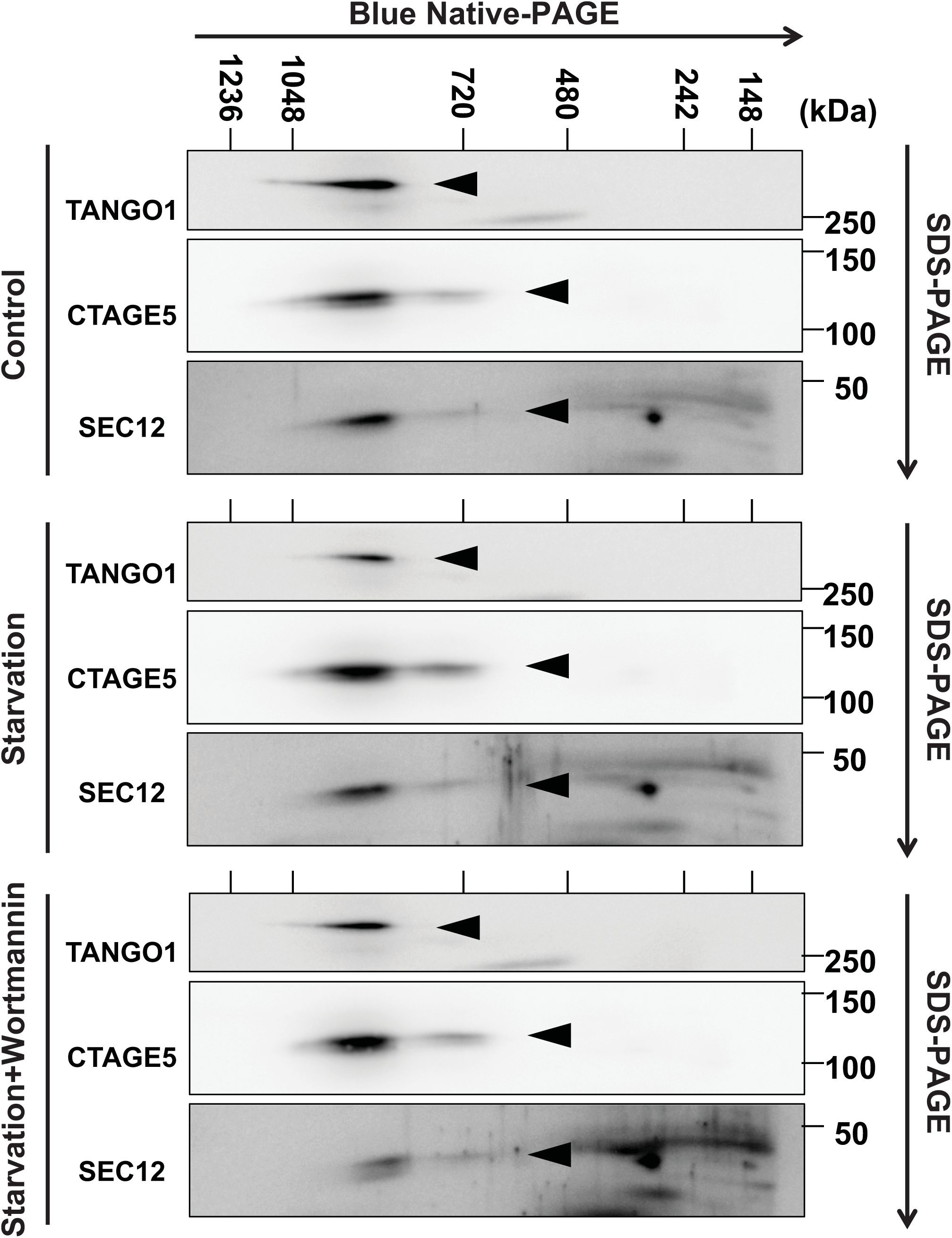
Blue Native-PAGE analysis of the TANGO1-CTAGE5-SEC12 complex. Hela cells were incubated in nutrient-rich medium or starved in EBSS for 1 h in the absence or presence of 20 nM wortmannin. Cells were lysed and Blue Native-PAGE was performed to separate protein complexes. SDS-PAGE and immunoblot were followed to determine the migration of the indicated proteins in the Blue Native-PAGE gel. Triangles point to the indicate protein bands.

**Figure 5.**
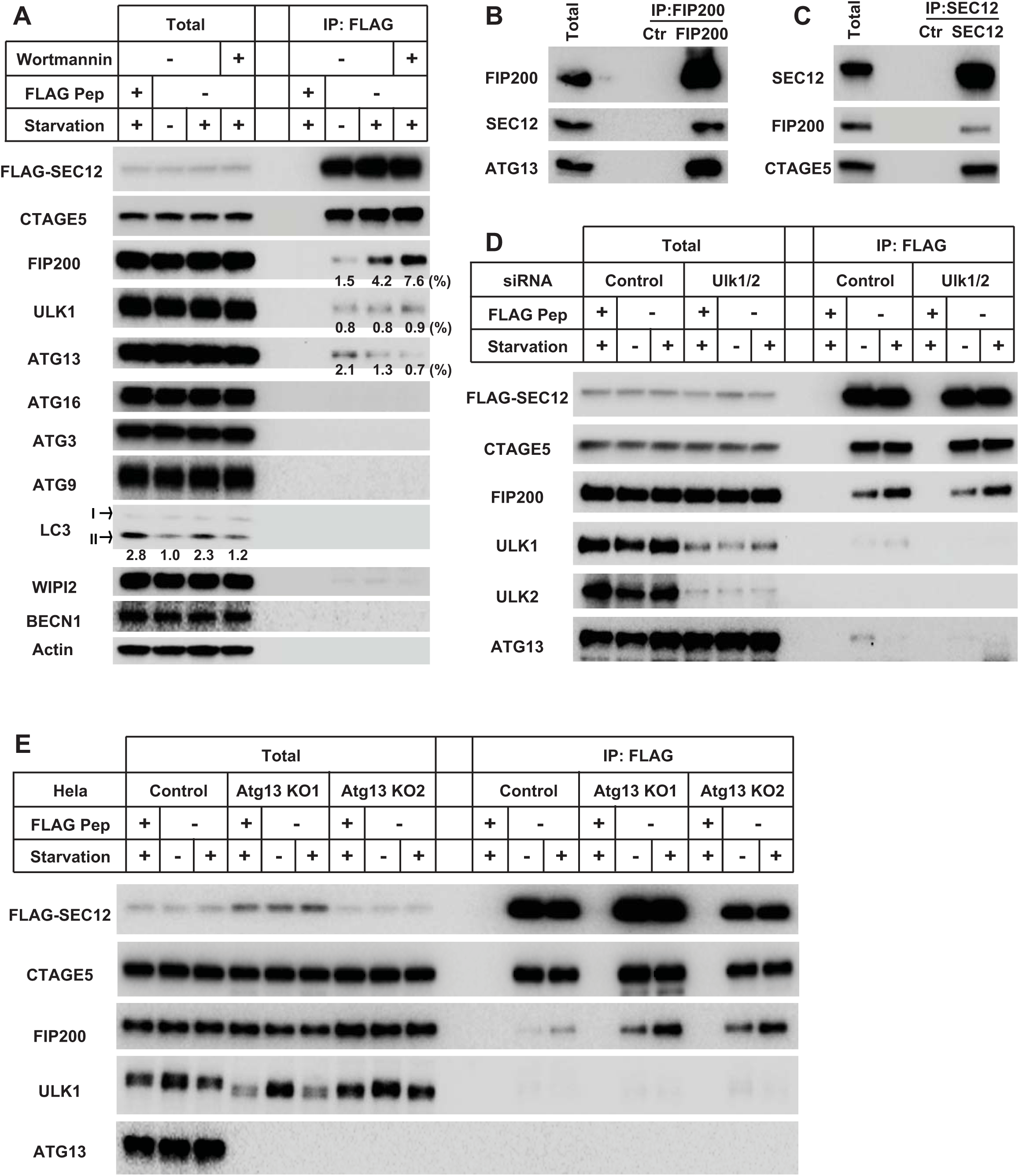
The association between SEC12 and FIP200 is independent of ULK1/2 and ATG13. (A) HEK293T cells were transfected with plasmids encoding FLAG-SEC12. After 24 h, the cells were incubated in nutrient-rich medium or starved in EBSS for 1 h in the absence or presence of 10 nM wortmannin. Immunoprecipitation of FLAG-SEC12 was performed and the levels of indicated proteins from indicated fractions were determined by immunoblot. Quantification of coIP shows the percentage of co-precipitated FIP200, ULK1, or ATG13 under the indicated conditions. Quantification of LC3 lipidation shows the relative level of LC3 lipidation to the control (before starvation). (B, C) Hela cells were starved in EBSS for 1 h. Immunoprecipitation of endogenous FIP200 (B) or SEC12 (C) was performed and the levels of indicated proteins from indicated fractions were determined by immunoblot. (D) HEK293T cells were transfected with control or siRNAs against Ulk1 and Ulk2. After 48 h, the cells were transfected with plasmids encoding FLAG-SEC12. After another 24 h, the cells were incubated in nutrient-rich medium or starved in EBSS for 1 h. Immunoprecipitation of FLAG-SEC12 was performed and the levels of indicated proteins from indicated fractions were determined by immunoblot. (E) Hela cells of control and Atg13 KO generated by CRISPR/Cas9 were transfected with plasmids encoding FLAG-SEC12. After 24 h, the cells were incubated in nutrient-rich medium or starved in EBSS for 1 h. Immunoprecipitation of FLAG-SEC12 was performed and the levels of indicated proteins from indicated fractions were determined by immunoblot.

Some ATG proteins, such as the mammalian ATG14 [47], yeast Atg17 (yeast functional homologue of FIP200)[48], yeast Atg16 [49], and lipidated yeast Atg8 or mammalian LC3/GABARAP [50,51], have been proposed to remodel autophagic membranes by tethering vesicles. Because starvation induced the enlargement of SEC12-ERES and also activated these ATG proteins, we considered the possibility that ATG proteins participated in the remodeling of SEC12-ERES. We used siRNA-mediated KD to deplete ATG proteins functioning in the early stages of autophagy including components of the ULK kinase complex (ULK1/2, FIP200 and ATG13) and PI3K complex (ATG14, BECN1). LC3 lipidation was compromised in the cells deficient in these ATG proteins (Fig 4A), confirming the effect of KD. FIP200 depletion abolished the enlargement of SEC12-ERES in starved cells (Fig 4B, C). Although FIP200 has been reported to form a complex with ULK1/2 and ATG13 [45,52], KD of ULK1/2 or ATG13 did not affect the remodeling of SEC12-ERES (Fig 4B, C), nor did the depletion of ATG14 or BECN1(Fig 4B, C), consistent with a similar minimum effect of inhibitors of PI3K (Fig EV2). Depletion of ATG16 or ATG5 had no effect on the remodeling of SEC12-ERES (data not shown), indicating that ATG16 and lipidation are not involved in the remodeling of SEC12-ERES.

We next determined the role of FIP200 in SEC12 relocation to the ERGIC. Depletion of ATG14 or treatment of cells with one of three VPS34-specific inhibitors largely reduced the localization of SEC12 on the isolated ERGIC (Fig 4D, E), which is consistent with our previous observation on the requirement of PI3K for the relocalization of SEC12 [25]. In line with the requirement of FIP200 for SEC12-ERES remodeling, depletion of FIP200 compromised the relocation of SEC12 to the ERGIC in starved cells (Fig 4D). Therefore, the data indicate that FIP200 and the autophagic PI3K act in different steps in the relocation of SEC12 wherein FIP200 may be involved in the remodeling of SEC12-ERES whereas PI3K may act later.

### FIP200 associates with SEC12 independent of ULK1/2 and ATG13

To test the possibility of SEC12-FIP200 association, we performed co-IP analysis. A small amount of FIP200 (^~^1.5%) co-precipitated with SEC12 under nutrient-rich conditions (Fig 5A). Interestingly, starvation enhanced the association between SEC12 and FIP200 (Fig 5A, ^~^3 fold increase). The PI3K inhibitor did not inhibit but moderately increased the formation of SEC12-FIP200 complex (Fig 5A, ^~^2 fold further increase), which was consistent with its positive effect on SEC12-ERES remodeling (Fig EV2). ULK1 weakly associated with SEC12 in all these conditions (^~^0.8%) whereas a marginal amount of ATG13 (^~^2.1%) co-precipitated with a response to starvation and PI3K inhibition opposite that of FIP200 (Fig 5A, ^~^1.5 fold decrease upon starvation and another ^~^1.5 fold decrease in the presence of wortmannin). CTAGE5 formed a complex with SEC12 irrespective of these treatments. The other ATG proteins we examined (ATG16, ATG3, ATG9, LC3, WIPI2 and BECN1) did not associate with SEC12 (Fig 5A). Endogenous FIP200 and SEC12 also associated with each other in the co-IP analysis (Fig 5B, C)

Although FIP200 forms a complex with ULK1/2, ATG13 and ATG101 in the initiation of PAS formation [45,52,53], the sole requirement of FIP200 in SEC12-ERES remodeling (Fig 4) and the unique starvation-enhanced FIP200-SEC12 complex formation versus ULK1 and ATG13 (Fig 5A) indicated that FIP200 may act independently of the other ULK kinase complex components. To test this further, we performed co-IP in cells depleted of ULK1/2 by siRNA (Fig 5D). The starvation-enhanced association of FIP200 and SEC12 was similar in both control and ULK1/2 KD cells (Fig 5D). We then performed a similar co-IP analysis in cells depleted of ATG13 by CRISPR/CAS9 (Fig 5E). Here the association between FIP200 and SEC12 was increased in ATG13-deficient cells and enhanced by starvation (Fig 5E). Therefore, these data indicate that FIP200 forms a complex with SEC12 which is enhanced by starvation and is independent of ULK1/2 and ATG13.

### FIP200 is enriched in the ERES/ERGIC region after starvation

The association between SEC12 and FIP200 indicates a role for FIP200 on the ERES. Consistent with a previous study [45], we found by confocal microscopy that in starved cells FIP200 formed punctate structures consistent with a PAS (Fig 6A, arrows pointed). Punctate FIP200 colocalized with WIPI2, another PAS marker [54] (Fig 6A, arrows), but showed less overlap with SEC12 (Fig 6A, B, arrows). FIP200 was also enriched at the perinuclear region positive for the SEC12-ERES and the ERGIC but not the PAS marker WIPI2 (Fig 6A, B, the big arrow head pointed region and the magnified view), suggesting that the pool of FIP200 associating with SEC12 may localize on the ERES/ERGIC.

**Figure 6.**
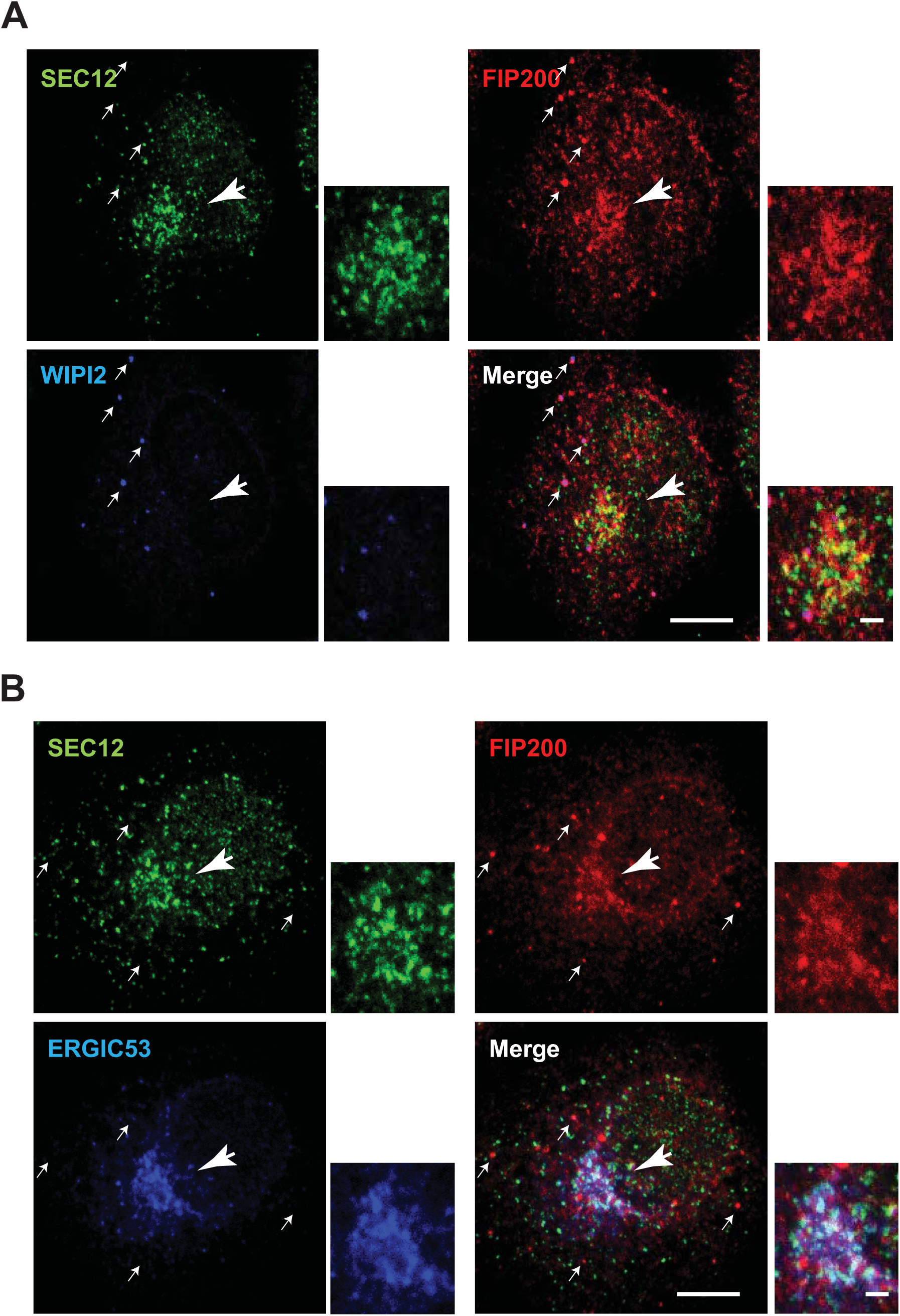
Enrichment of FIP200 around the ERES/ERGIC upon starvation. Hela cells were starved for 1 h. Immunofluorescence was performed to analyze the localization of SEC12, FIP200 and WIPI2 (A), and SEC12, FIP200 and ERGIC53 (B). Bar: 10 μm (2 μm in the magnified view)

### The C-terminus of FIP200 is required for association with SEC12, SEC12-ERES remodeling and autophagosome biogenesis

To probe the domain structure required for FIP200-SEC12 association, we truncated FIP200 into three fragments (N, N-terminus, M, Middle, and C, C-terminus) (Fig 7A) [55] and performed co-IP analysis (Fig 7A). The N-terminus of FIP200 associated with the rest of the ULK kinase complex, ULK1, ATG13 and ATG101 (Fig 7A). Although both the N-terminus and middle fragment of FIP200 were detected in association with SEC12, the C-terminus appeared to interact more robustly (Fig 7A). We also mapped the region on SEC12 that associates with FIP200 (Fig EV5). As shown in Fig EV5, the C-terminal half of the SEC12 cytoplasmic domain (corresponding to aa240-386) forms a complex with FIP200 (Fig EV5).

**Figure 7.**
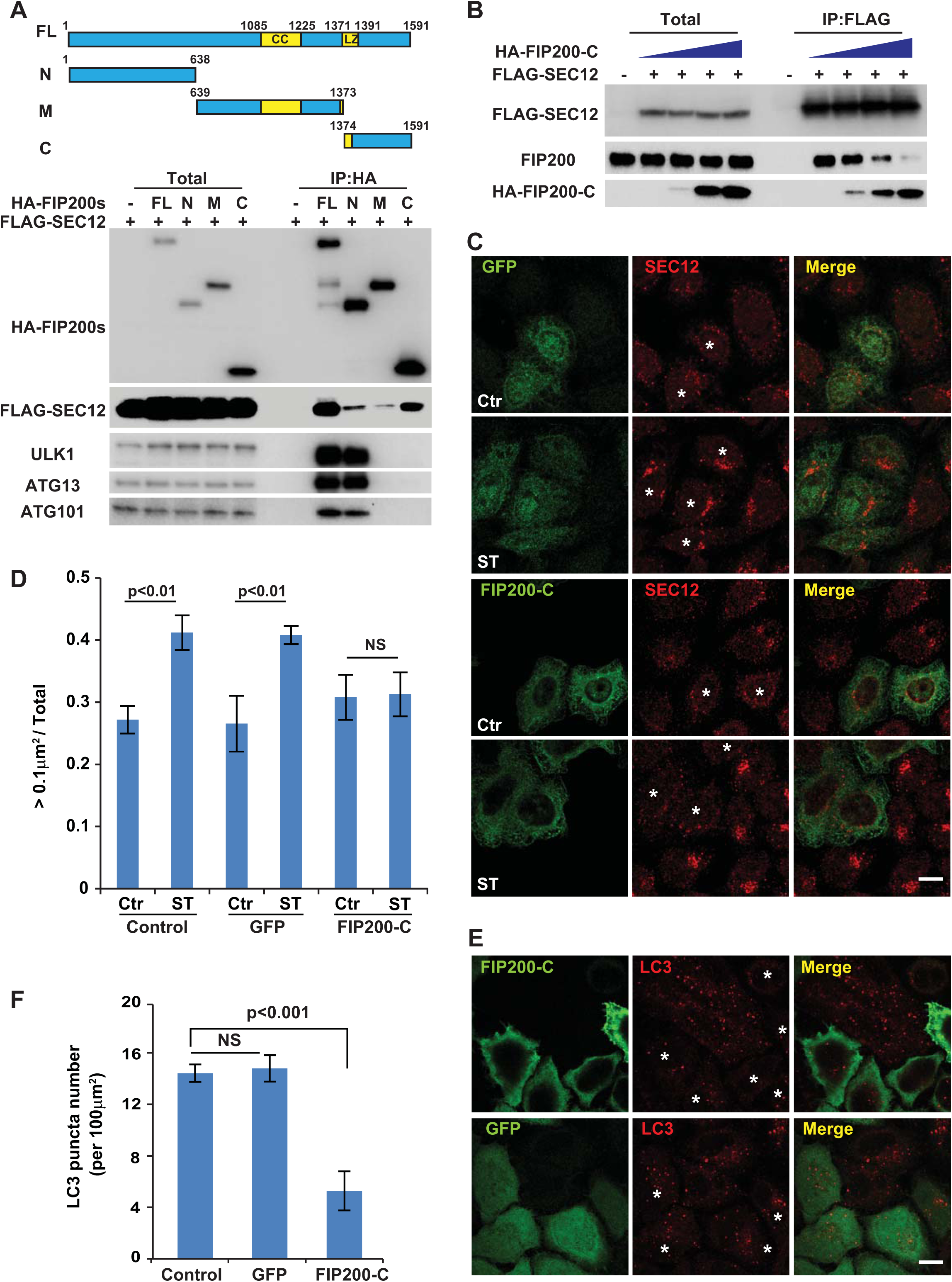
The association between SEC12 and C-terminal domain of FIP200 is required for the remodeling of SEC12-ERES and autophagosome biogenesis. (A) HEK293T cells were transfected with plasmids encoding FLAG-SEC12 or together with plasmids encoding HA-FIP200 variants. After 24 h, the cells were starved in EBSS for 1 h. Immunoprecipitation of HA-FIP200 was performed and the levels of indicated proteins from indicated fractions were determined by immunoblot. Schematic view of the FIP200 truncations is shown in the top. (B) HEK293T cells were transfected with increasing amounts of plasmids encoding HA-FIP200 C-terminus together with plasmids encoding FLAG-SEC12. After 24 h, the cells were starved in EBSS for 1 h. Immunoprecipitation of FLAG-SEC12 was performed and the levels of indicated proteins from indicated fractions were determined by immunoblot. (C) Hela cells were transfected with plasmids encoding GFP or the HA-FIP200 C-terminus. After 24 h, the cells were incubated in nutrient-rich medium or starved in EBSS for 1 h. Immunofluorescence and confocal microscopy were performed to visualize SEC12 (asterisks mark the cells with GFP or FIP200-C), GFP and HA-FIP200 C-terminus. Bar: 10 μm (D) Quantification of the fraction of SEC12 compartments larger than 0.1 μm^2^ in area analyzed in (C). Error bars represent standard deviations. P value was obtained from Two-tailed T test. Five experiments (50-100 cell/experiment) were performed for the statistics. Control: cells without expression of GFP or HA-FIP200 C-terminus; GFP: cells with GFP expression; FIP200-C: cells with HA-FIP200 C-terminus expression. (E) Hela cells were transfected with plasmids encoding GFP or the HA-FIP200 C-terminus. After 24 h, the cells were starved in EBSS for 1 h. Immunofluorescence and confocal microscopy were performed to visualize LC3 (asterisks mark the cells with GFP or FIP200-C), GFP and HA-FIP200 C-terminus. Bar: 10 μm (F) Quantification of the LC3 puncta number analyzed in (E). Error bars represent standard deviations. P value was obtained from Two-tailed T test. Five experiments (50-100 cell/experiment) were performed for the statistics. Control: cells without expression of GFP or HA-FIP200 C-terminus; GFP: cells with GFP expression; FIP200-C: cells with HA-FIP200 C-terminus expression.

**Expanded View Figure 5.**
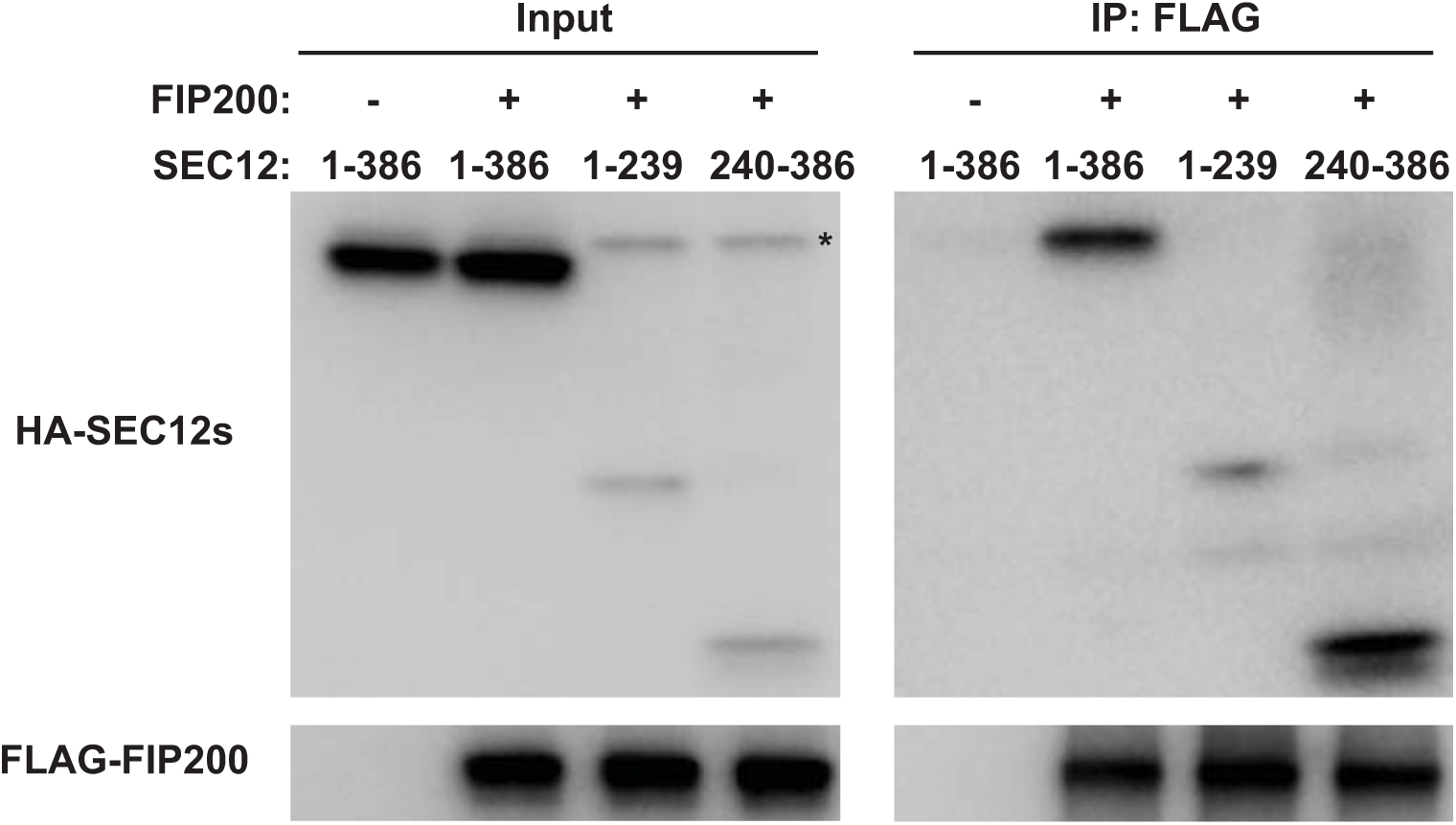
The association between SEC12 cytoplasmic domains and FIP200. HEK293T cells were transfected with plasmids encoding SEC12 cytoplasmic domain variants together with FLAG-FIP200. After 24 h, the cells were starved in EBSS for 1 h. Immunoprecipitation of HA-FIP200 was performed and the levels of indicated proteins from indicated fractions were determined by immunoblot. Asterisk indicates a non-specific band.

Because the major SEC12 association region in FIP200 differed from the domain apparently responsible for its interaction with the ULK kinase complex, we sought to determine the specific role of FIP200-SEC12 association in autophagosome biogenesis in cells by overexpressing the C-terminal fragment of FIP200. Indeed, exogenous expression of the C-terminal FIP200 blocked endogenous FIP200 association with SEC12 (Fig 7B). We then determined the effect of C-terminal FIP200 expression on the starvation-induced remodeling of SEC12-ERES. As shown in Fig 7C, D, starvation increased the size of SEC12-ERES in control cells not expressing exogenous protein or in cells expressing GFP alone (Fig 7C, D, GFP cells are marked by asterisks). In the presence of the FIP200 C-terminus, the enlargement of SEC12-ERES was inhibited (Fig 7C, D, compare cells expressing FIP200-C marked by asterisks with surrounding control cells without FIP200-C). Furthermore, the overexpression of FIP200 C-terminus blocked autophagosome biogenesis as indicated by a decreased number of LC3 puncta in starved versus control cells, whereas GFP expression had no effect (Fig 7E, F, compare cells expressing FIP200-C marked by asterisks with surrounding control cells without FIP200-C). Therefore, FIP200 has another role in autophagosome biogenesis: in addition to a function in the ULK kinase complex, it also associates with SEC12 to facilitate the starvation-induced remodeling of SEC12-ERES which is important for supplying membranes for autophagosome biogenesis.

## Discussion

COPII transport vesicle formation from the ER and the ERGIC has been linked to the process of autophagosome biogenesis [25,32,56]. The ERES is the site from which COPII vesicles acquire regulated and constitutive secretory and membrane cargo proteins that are conveyed to the ERGIC [28]. Although the ERGIC then delivers vesicles or is transformed into the cis Golgi cisterna for the traffic of normal cargo proteins, we have found that a novel pool of COPII vesicles bud from the ERGIC in starved cells that are activated for a PI3K, and have suggested that these novel COPII vesicles may be the precursor of the phagophore membrane [25]. Consistent with this, a recent study using sophisticated imaging approaches found that the ERGIC but not the conventional ERES is associated with markers decorating the PAS [27]. In a previous report we demonstrated that a fraction of the pool of the membrane protein SEC12, the nucleotide exchange catalyst for SAR1 which upon activation initiates the assembly of the COPII coat, is transferred to the ERGIC in starved cells [25]. Using super-resolution imaging, we now report that a starvation-induced remodeling of ERES facilitates the relocation of SEC12 to the ERGIC (Fig 1) as a potential prelude for the assembly of ERGIC-COPII vesicles. ERES remodeling is dependent on CTAGE5 (Fig 2), a coiled-coil domain protein reported to directly bind to and concentrate SEC12 on the ERES [43]. Depletion of CTAGE5 abolishes the enrichment of SEC12 on the ERES as well as starvation-induced remodeling, thereby inhibiting COPII assembly on the ERGIC and thus autophagosome biogenesis (Fig 2, 3). Further, we found that the autophagic factor FIP200, a subunit of the ULK protein kinase complex required to initiate phagophore assembly [45], associates with SEC12 via its C-terminal domain independent of other subunits of the ULK complex (e.g. ATG13 and ULK1/2) and facilitates the remodeling of SEC12-ERES (Fig 4-7).

As the initiator of COPII vesicle formation, the concentration of SEC12 on the ERES is regulated to accommodate demands for cargo packaging. For example, in the event of large cargo secretion (collagen VII), SEC12 concentration on the ERES via association with CTAGE5 is required for large cargo export out of the ER [43]. Concentration of SEC12 may facilitate the efficient activation of SAR1 to enhance the packaging of large cargoes [57]. Our study further suggests that the concentration of SEC12 on the ERES facilitates its relocation to the ERGIC to trigger the assembly of ERGIC-COPII vesicles and thus autophagosome biogenesis (Fig 2, 3).

Starvation appears to result in a change in the size and shape of the SEC12-ERES (Fig.1). The morphologic change appears dependent on a starvation-induced association between SEC12 and FIP200 (Fig 5). FIP200 is known to form a complex with ULK1/2, ATG13 and ATG101 to nucleate the assembly of the PAS as an initial step in the formation of the autophagic membrane. We find that FIP200 plays a special role in starvation-induced remodeling of SEC12-ERES independent of the other components in the complex. FIP200, but not ULK1/2 or ATG13, is required for the remodeling of SEC12-ERES (Fig 5). In contrast to FIP200, we find a less abundant and starvation-independent association of ULK1/2 and ATG13 with SEC12 (Fig 5). Further, the association between FIP200 and SEC12 is not dependent on ULK1/2 or ATG13 (Fig 5). These complexes are also distinguished by the interaction of SEC12 and C-terminal domain of FIP200 whereas an N-terminal domain of FIP200 appears to be the site of interaction with the ULK1 complex (Fig 7A). We thus suggest that FIP200 may act independent of the ULK complex to facilitate the starvation response that mobilizes a fraction of the pool of SEC12 from the ERES to the ERGIC.

These simple molecular characteristics of the complex do not illuminate the means by which starvation induces SEC12-ERES to become reorganized. Considering Atg17, the yeast functional homolog of FIP200, forms a homodimer that is proposed to scaffold for tethering vesicles [48,58], FIP200 may do likewise and cluster SEC12 molecules in the ERES, leading to the increase of membrane size. The effect of starvation on the oligomeric state of FIP200 bears investigation.

A recent study indicated an ATG13-independent role of ULK1 in the phosphorylation of SEC16A, which is thought to regulate the formation of ER-derived COPII vesicles [59]. Although SEC16A is required to maintain ERES organization, we found that ULK1 is dispensable for starvation-induced remodeling of the ERES (Fig 4). Therefore, the ULK1-mediated phosphorylation of SEC16A may not be involved in the remodeling of the ERES during starvation and the induction of autophagy.

Our 3D-STORM images probe the morphology of the enlarged ERES and its spatial relationship with the ERGIC in starved cells (Fig 1 and Appendix Fig S1 & S2). Such imaging has allowed us to directly document the relocation of SEC12 to the ERGIC although the molecular mechanism of this remains unclear. It is noteworthy that in the steady state, SEC12 is bound to CTAGE5 at the ERES, but SEC12 unbound to CTAGE5 is found in the ERGIC isolated from starved cells (Fig 2F, G). One possibility is that starvation may induce dissociation of a fraction of SEC12 molecules from a complex with CTAGE5 thus releasing it for transport to the ERGIC. We previously found that ^~^2% of the total pool of SEC12 was mobilized to the ERGIC in starved cells (data not shown). Such a small amount of SEC12 might be beyond the limit of detection in our co-IP assay where we saw no change in the complex of SEC12 and CTAGE5 with and without starvation of cells (Fig EV3A and Fig 5). Another possibility is that a pool of SEC12 unbound to CTAGE5 exits at the ERES and relocates to the ERGIC upon starvation. In this situation, CTAGE5 may act as a scaffold, like SEC16, to maintain the proper structure of the SEC12-ERES. The two possibilities remain to be explored.

Our previous study indicated a requirement for the autophagic PI3K in the relocation of COPII to the ERGIC [25]. However, here we find that PI3K is not required for the remodeling of the ERES (Fig EV2). At which step does PI3K facilitate the relocation of COPII? Studies in yeast cells indicate that SEC12 is retained in the ER by a selective cis Golgi-ER recycling pathway mediated by the RER1 protein [60-62]. We suggest that the starvation-induced activation of PI3K to produce PI3P on the ERGIC may serve to retain a fraction of SEC12 and of the COPII machinery at this location, possibly by interfering with the action of the RER1 protein.

In contrast to mammalian cells, the yeast (*Saccharomyces cerevisiae*) lacks a distinct ERGIC structure. In place of the ERGIC, specialized COPII vesicles may bud directly from the ERES to supply the autophagic membrane. According to a recent study, phosphorylation of Sec24, the cargo receptor subunit of COPII, may be a mechanism for reprogramming the COPII vesicles towards autophagic membrane formation in yeast [63].

Our current study, together with the previous ones [23,25], suggests a model for a remodeling of the endomembrane system, facilitated by autophagic factors (FIP200, PI3K), scaffold proteins (CTAGE5), and membrane remodeling proteins (COPII)-(Fig 8). Starvation induces CTAGE5- and FIP200-dependent remodeling of the SEC12-ERES leading to the PI3K-dependent relocation of SEC12 to the ERGIC (Fig 8A, B). Although this bears more investigation, FIP200 may associate with SEC12 on the ERES to facilitate the remodeling. ERGIC-localized SEC12 triggers the assembly of ERGIC-COPII vesicles as membrane templates for LC3 lipidation (Fig 8C).

**Figure 8.**
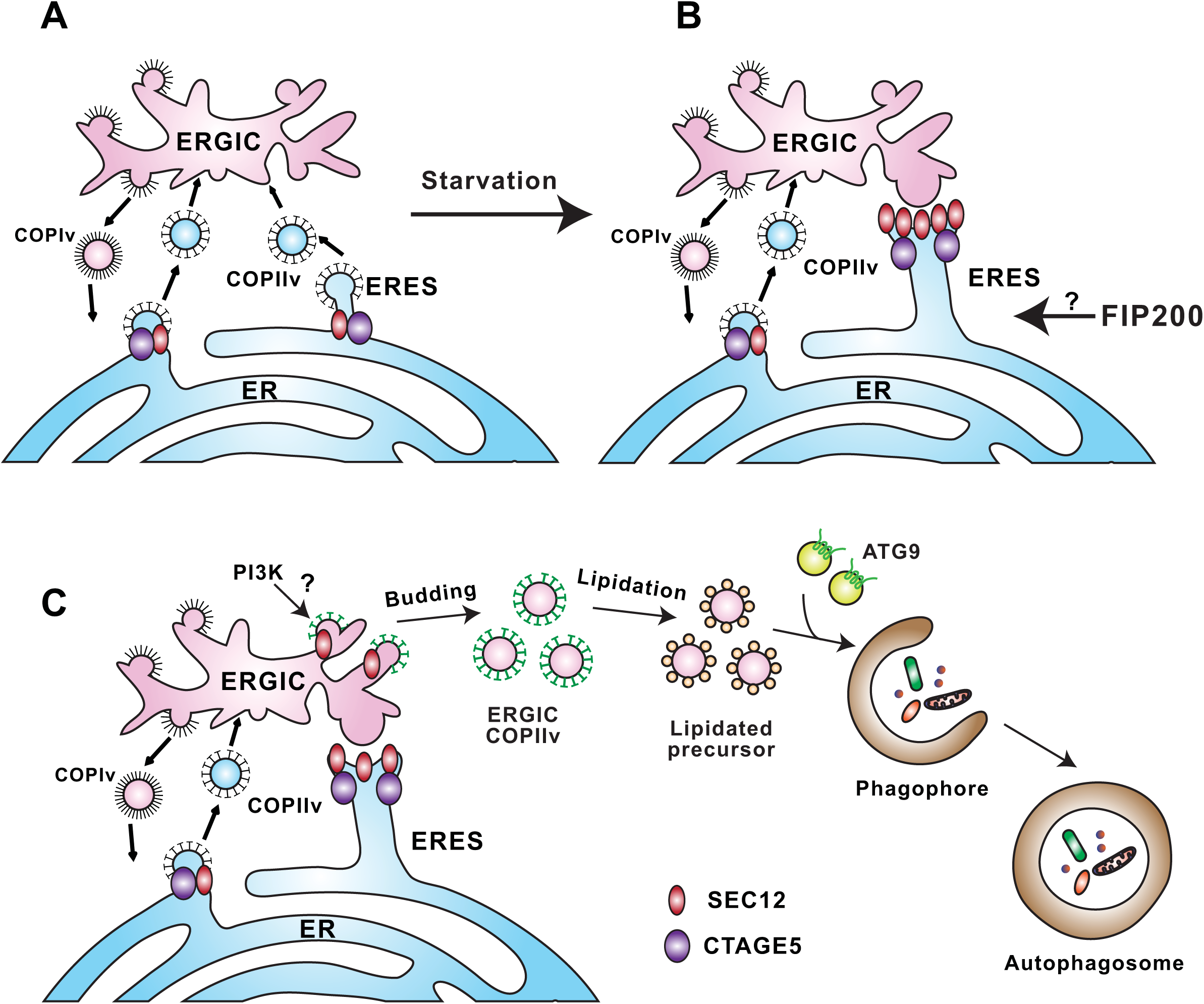
A proposed model for remodeling of the SEC12-ERES leading to the generation of autophagosomal precursors. (A) In steady state conditions, SEC12 is enriched in the ERES dependent on association with CTAGE5 for protein cargo transport by packaging COPII vesicles. (B) Upon starvation, SEC12-ERES is enlarged and surrounds the ERGIC, dependent on FIP200 and CTAGE5. (C) The remodeling of SEC12-ERES leads to the relocation of SEC12 to the ERGIC (dependent on the autophagic PI3K) to trigger the assembly of ERGIC-COPII vesicles as a membrane template for LC3 lipidation, a potential vesicular pool for the assembly of the pre-autophagosomal membrane.

ATG9-positive vesicles have been shown to traffic from multiple membrane compartments including the Golgi [64,65], endosomes [34,66], and certain intermediate compartments [67] to deliver membranes to the PAS [68,69]. Recent evidence indicates that markers of the PAS individually associate with the ERGIC and ATG9, indicating two separate pools of autophagic membrane sources [58]. In addition, the two membrane sources seem to interact based on at least two pieces of evidence: 1) ERGIC and ATG9 compartments are localized closely [58] and 2) COPII machinery associates with ATG9 [63]. Therefore, it is possible that the autophagic COPII vesicles may coalesce with a distinct pool of ATG9 vesicles or fuse homotypically to form the pre-autophagosome (Fig 8C).

## Materials and Methods

### Materials, antibodies, plasmids and cell culture

Wortmannin, 3-methyladenine and bafilomycin A1 were as previously described [23,25]. We purchased PIK-III and SAR405 from APExBIO (Houston, TX), and VPS34-IN1 from Cayman (Ann Arbor, MI). Control siRNA was as described previously [23] and other siRNAs were purchase from Qiagen (Germantown, MD). SiRNAs used were human cTAGE5 (CAGGATATTATCTAATGGTGA, ACCAATAACTTTGAGGTGCAA, AGGGCAGATTATTTCCCATGA, TAGACGGTATAACTATTTCAA), Ulk1 (CGCGCGGTACCTCCAGAGCAA, TGCCCTTTGCGTTATATTGTA), Ulk2 (CAGCCTGAGATACGTGCCTTA, CAAGCATTTGATCGACCCAAA, AAGACTTGGGAGACATGTTAA, TCGCAGATTATTTGCAAGCGA), FIP200 (CTGGGACGGATACAAATCCAA, ACGCAAATCAGTTGATGATTA), Atg13 (TCGCGGAGTCTTAGGAGCAAA, CTGGGCGTATGATTGACTTAA, CTCCGTAATGAGAGCCCGGAA, AAGGCGGGAGTGACCGCTTAA), Atg14 (CTCGGTGACCTCCTGGTTTAA, TTGGATTAGCCTCCCTAACAA, CTGCATACCCTCAGGAATCTA, CCGGGAGAGGTTTATCGACAA) and Beclin-1 (GAGGATGACAGTGAACAGTTA, TGGACAGTTTGGCACAATCAA, AGGGTCTAAGACGTCCAACAA, ACCGACTTGTTCCTTACGGAA). The transfection of the siRNA was performed as previously described (Ge 2014).

We purchased mouse anti-ERGIC53 (G1/93) from Enzo Life Sciences (Farmingdale, NY, G1/93 No.ALX-804-602-C100), rabbit anti-ULK1 and ATG101 from Cell Signaling (Danvers, MA, No.8674), rabbit anti-ULK2 from Thermo Fisher (Waltham, MA, No.PA5-22173), mouse anti-ATG13, anti-ATG16, and rabbit anti-P62 from MBL (Woburn, MA, No. PM045), mouse anti-WIPI2 from AbD Serotec (Raleigh, NC, No.MCA5780GA), rabbit anti-FIP200 for immunofluorescence and immunoprecipitation from Proteintech (Rosemont, IL, No.17250-1-AP), goat anti-SEC12 for immunoprecipitation from R&D Systems (Minneapolis, MN, No.AF5557). Rabbit anti-CTAGE5 and rat anti-SEC12 were as previously described [43]. SEC22B, FIP200, ATG14, Beclin-1, LC3, ATG9, HA and FLAG antibodies were described before [23,25].

The plasmid encoding the HA-FIP200 full length was obtained from addgene [52]. The plasmids encoding the HA-FIP200 truncations were constructed by PCR amplification of the corresponding regions described before [55]. The amplified fragments were inserted into the same backbone vector of HA-FIP200 full length. N-terminal HA-tagged SEC12 domains (1-386, 1-239, and 240–386 aa) were cloned into pCMV5 vectors that were gifts from David Russell (University of Texas Southwestern Medical Center).

Other reagents were described previously [23,25].

### Cell culture and transfection

Maintenance of cell lines and transfection were described previously [23,25]. The Hela and HEK293T cell lines (originally from ATCC) were obtained from the Berkeley tissue culture facility. And they have been tested by the facility to be clear of mycoplasma.

### Generation of CRISPR/CAS9 KO cell lines

Hela or HEK293T cells were transfected with a pX330 vector-derived plasmid [70] containing the targeting sequence from cTAGE5 (ACATTCTCTTAGTATAGCAC) or Atg13 (GAATGGACACATTACCTTGA), and a PGK promotor-driven Venus construct (reconstructed by Liangqi Xie from Robert Tjian lab at UC Berkeley). After a 24 h transfection, FACS sorting was performed to inoculate single transfected cells in each well of 96-well plates. After two weeks, single colonies were expanded and validated by immunoblot and DNA sequencing of the targeted area. Validated positive colonies were employed for the experiments. The cell lines were also validated to be clear of mycoplasma.

### Blue Native-PAGE

HeLa cells were either untreated or starved with EBSS in the absence or presence of 20 nM wortmannin for 1 h before extraction. Blue Native-PAGE analysis was performed essentially as described previously [71]. Cells were extracted with BN buffer (20 mM Bis-Tris, 500 mM ε-amino capronic acid, pH 7.0, 20 mM NaCl, 2 mM EDTA, 10 % glycerol, and protease inhibitors) containing 1 % digitonin and centrifuged at 20,000 g at 4°C The cell lysates were supplemented with final 0.25 % of CBB G-250 for electrophoresis. In the first dimension of Blue Native-PAGE, 4-15% gradient gel was run at 4°C with CBB+ cathode buffer (50 mM Tricine, 15 mM Bis-Tris, PH 7.0, and 0.02% CBB G-250) and anode buffer (50 mM Bis-Tris, pH 7.0). The CBB+ cathode buffer was exchanged with CBB-cathode buffer (50 mM Tricine, 15 mM Bis-Tris, PH 7.0) once the dye front migration reached one third of the gel. For further separation in a second-dimension SDS-PAGE, we cut the gel lanes and heated to 100 °C in Laemmli Sample buffer. The gel strip was washed with SDS-PAGE buffer (25 mM Tris, 192 mM glycine, PH8.3 and 0.1 % SDS) and placed on the stacking part of an SDS-PAGE gel. The second dimension SDS-PAGE was electrophoresed in SDS-PAGE buffer at room temperature.

### Co-immunoprecipitation

The cells were treated with indicated conditions, harvested and washed once with PBS. The cell pellets from one 10 cm dish were lysed with 1 ml co-IP buffer (20 mM Tris-HCl, PH7.5, 150 mM NaCl, 1 mM EDTA and 0.5% NP-40) by passing samples through 22G needles. The lysates were centrifuged at 20,000x g for 15 min in a microfuge at 4 °C. The supernatant fractions were transferred to tubes and incubated with 40 μl (1:1 slurry) anti-FLAG agarose (Sigma, St. Louis, MO) in the absence or presence of 0.02 mg/ml 3XFLAG peptides (David King, UC Berkeley) for 3 h at 4 °C. For coIP to determine endogenous protein association, 5μg antibodies was added to the supernatant and incubated for 2 h at 4 °C. 40 μl (1:1 slurry) protein A/G agarose was then added and incubated for another 1 h at 4 °C. The agarose in each sample was washed 4 times with 1 ml co-IP buffer. Proteins bound to the agarose were eluted with 40 μl 1 mg/ml 3XFLAG peptides at room temperature for 40 min (FLAG IP)or eluted with 100 μl sample loading buffer (endogenous protein IP).

### Membrane fractionation and immunoblot

These were performed as previously described [23,25,72,73]. Quantification of SEC12 relocation to the ERGIC was based on the percentage of ERGIC SEC12 relative to total SEC12. Quantification of LC3 lipidation was based on the ratio of LC3-II to actin normalized to control treatment in nutrient-rich conditions. Quantification of FIP200, ATG13, and ULK1 in the coIP experiment was based on the percentage of FIP200, ATG13, or ULK1 in the pellet fraction relative to the total protein in the input fraction.

### Immunofluorescence microscopy and quantification

Immunofluorescence was performed as previously described [72,73]. Confocal images were acquired with a Zeiss LSM 710 laser confocal scanning microscope (Molecular Imaging Center, UC Berkeley). Colocalization of the confocal images was calculated by a pixel-based method using Image J with RGB Profiler plugin. SIM images were collected using the Elyra PS.1 microscope (Carl Zeiss Microscopy). A 3D surface model was generated and quantification of the volume of SEC12-ERES was carried out using Imaris 7.7.1 software (CNR, Biological Imaging Facility, UC Berkeley). Quantification of the area of SEC12-ERES, CTAGE5, and SEC16 puncta was performed using the Analyze Particles function of Image J as described previously [25]. We chose 0.1 μm^2^/0.04 μm^3^as the cutoff for quantification because in STORM images it was the lower size limit of the SEC12 structure that remodeled after starvation. The images were collected unbiasedly and under optimized settings to avoid signal saturation. Quantification of the number of FIP200 and LC3 puncta was performed with a similar approach using Image J [25].

### 3D-STORM microscopy

Dye-labeled cell samples were mounted on glass slides with a standard STORM imaging buffer consisting of 5% (w/v) glucose, 100 mM cysteamine, 0.8 mg/ml glucose oxidase, and 40 μg/ml catalase in Tris-HCl (pH 7.5) [41,42]. Coverslips were sealed using Cytoseal 60. STORM imaging was performed on a homebuilt setup based on a modified Nikon Eclipse Ti-U inverted fluorescence microscope using a Nikon CFI Plan Apo λ 100x oil immersion objective (NA 1.45). Dye molecules were photoswitched to the dark state and imaged using either 647- or 560-nm lasers (MPB Communications); these lasers were passed through an acousto-optic tunable filter and introduced through an optical fiber into the back focal plane of the microscope and onto the sample at intensities of ^~^2 kW cm^-2^. A translation stage was used to shift the laser beams towards the edge of the objective so that light reached the sample at incident angles slightly smaller than the critical angle of the glass-water interface. A 405-nm laser was used concurrently with either the 647- or 560-nm lasers to reactivate fluorophores into the emitting state. The power of the 405-nm laser (typical range 0-1 W cm-2) was adjusted during image acquisition so that at any given instant, only a small, optically resolvable fraction of the fluorophores in the sample were in the emitting state. For 3D STORM imaging, a cylindrical lens was inserted into the imaging path so that images of single molecules were elongated in opposite directions for molecules on the proximal and distal sides of the focal plane [42]. Data was collected at a framerate of 110 Hz using an Andor iXon Ultra 897 EM-CCD camera, for a total of ^~^80,000 frames per image. The raw STORM data was analyzed according to previously described methods [41,42].

Three-color imaging was performed on targets labeled by Alexa Fluor 647, CF680, and CF568 via sequential imaging with 647-nm and 560-nm excitation. With 647-nm excitation, a ratiometric detection scheme [74,75] was employed to first concurrently collect the emission of single Alexa Fluor 647 and CF680 molecules. Emission of single molecules was split into two light paths (channels) using a long pass dichroic mirror (T685lpxr; Chroma), each of which were projected onto one half of an Andor iXon Ultra 897 EM-CCD camera. Fluorophore assignment was performed by localizing and recording the intensity of each single molecule in the two channels. Excitation at 560 nm was subsequently used to image CF568 through the reflected light path of the dichroic mirror.

## Acknowledgement

We thank David Melville and Steven Ruzin (UC Berkeley) for technical assistance on image acquisition and quantification; Robert Tjian, Liangqi Xie and David King (UC Berkeley) for reagents; Bob Lesch, Alison Killilea and Carissa Tasto (UC Berkeley) for lab service and tissue culture; Kartoosh Heydari (UC Berkeley) for cell sorting; James Hurley (UC Berkeley) for helpful discussion. L.G. is supported by NIH Pathway to Independence Award (Parent K99/R00) National Institute of General Medical Sciences (Grant Number: 1K99GM114397-02). M.Z. is an HHMI Associate. S.K. and K.X. acknowledge support from NSF under CHE-1554717, the Pew Biomedical Scholars Award, and the Sloan Research Fellowship. A.M. and D.L. are undergraduates at UC Berkeley. R.S. is an Investigator of the HHMI and a Senior Fellow of the UC Berkeley Miller Institute. M.M. and K.S. are members of Toshiaki Katada lab supported by Japan Society for the Promotion of Science (JSPS, K.S.).

## Author contributions

L.G., K.X., and R.S. designed the experiments and study. L.G., M.Z., S.K., D.L., M.M., K.S., and A.M. carried out research experiments. All authors analyzed data. L.G., K.S., K.X., and R.S. wrote the paper.

